# Identification of pre-existing ubiquitous neoantigen-reactive tumor-infiltrating T-cells in a patient with metastatic pancreatic neuroendocrine tumor

**DOI:** 10.64898/2026.07.30.741721

**Authors:** Jean-Benoit Tanis, Katy McCann, Francisco Emmanuel Castañeda-Castro, Jaya Thomas, Alistair Bailey, Prabhat Singh, Eve Currall, Lindsey Chudley, Hayley Simon, Benjamin Nicholas, Judith Cave, Arjun Takhar, Susanne Burdak-Rothkamm, Stephen P. Schoenberger, Jason Greenbaum, Paul Skipp, Pandurangan Vijayanand, Gregory Seumois, Natalia Savelyeva, Christian Ottensmeier

**Author notes:** J-BT and KM contributed equally to this work. **Corresponding Author:** Prof Christian H. Ottensmeier, Professor of Immuno-Oncology, Liverpool Head and Neck Center, Institute of Systems, Molecular and Integrative Biology, University of Liverpool, Liverpool, L69 7ZX, UK.

## Abstract

**Background:** Mutation-derived neoantigens, typically identified in primary tumors, are emerging therapeutic targets for personalized cancer vaccines and adoptive T-cell therapies. However, clinical efficacy of neoantigen-directed therapies in patients with metastatic disease remains limited, partly due to inter-site genetic heterogeneity. We investigated whether ubiquitous neoantigens–derived from mutations shared across all tumor sites–could provide more effective, durable targets, particularly in patients undergoing resection of metastatic lesions.

**Methods:** Whole-exome and RNA sequencing were performed on 14 tumor samples (primary and 13 synchronous nodal metastases) from a treatment-naïve patient with pancreatic neuroendocrine tumor (PNET). Ubiquitous mutations were identified bioinformatically, and their immunogenicity assessed using in-vitro stimulation of autologous peripheral blood mononuclear cells followed by IFN-γ ELISpot assay. Neoantigen-specific T-cell clonotypes were further identified by HLA-tetramer staining and single-cell RNA/TCR sequencing. Neoantigen-reactive clonotypes identified in peripheral blood were tracked across multiple metastatic sites using bulk TCRβ repertoire sequencing.

**Results:** Among 1,195 non-synonymous mutations detected, eight were shared across all 14 tumor sites. Of these, one encoded a neoantigen that elicited a reproducible IFN-γ ELISpot response in peripheral blood, confirming its immunogenicity. Further, we identified the corresponding neoantigen-reactive TCR clonotypes in blood. Comparison with bulk TCRβ repertoires from eight metastatic sites showed that these clonotypes were present in every site analyzed, with evidence of local clonal expansion.

**Conclusion:** This study provides direct evidence that a single ubiquitous mutation-derived neoantigen can generate systemic T-cell responses and clonotype expansion across multiple metastatic sites in a TMB-low, TIL-low tumor. Our findings support incorporating mutation-sharing status across metastases as a key criterion for neoantigen selection in cancer vaccines and adoptive T-cell therapies. This approach could inform the design of neoantigen-directed immunotherapies in metastatic PNET and potentially other metastatic solid tumors.

**What is already known on this topic:** Neoantigen-directed therapies, such as personalized cancer vaccines or adoptive T-cell transfer, can induce anti-tumor responses but have shown limited success in metastatic disease. One major barrier is genetic heterogeneity between tumor sites, suggesting that targeting ubiquitous mutations–those shared across all tumor sites–may improve the efficacy of such therapies.

**What this study adds:** In one patient with metastatic pancreatic neuroendocrine tumor involving 13 lymph nodes, we identified eight ubiquitous mutations, one of which generated a detectable neoantigen-specific T-cell response in blood. The corresponding T-cell clonotypes were found across all metastatic sites analyzed and showed evidence of clonal expansion, providing direct evidence of systemic and local recognition of a shared neoantigen in a TMB-low/TIL-low cancer.

**How this study might affect research, practice or policy:** These findings support incorporating mutation sharing across metastases as a key criterion in neoantigen selection for cancer vaccines and adoptive T-cell therapies. This strategy could enhance the relevance and durability of neoantigen-directed approaches in patients with metastatic disease.

## BACKGROUND

Cancer immunotherapy has revolutionized treatment for many malignancies, yet challenges persist in metastatic disease^(1, 2)^. Neoantigens have emerged as promising targets for cancer immunotherapy ^(3, 4, 5)^. Clinical trials of neoantigen-directed therapy (NDT), such as personalized cancer vaccines, have demonstrated clinical benefit in the adjuvant setting^(6, 7)^, but limited success in advanced metastatic disease^(8, 9, 10)^.

A critical barrier to effective NDT is the heterogeneity between the primary tumor and its associated metastases, with mutation sharing across tumor sites ranging between 25-90% depending on cancer type and mode of metastases^(11, 12, 13)^. This disparity is more pronounced in extensive metastatic disease, where as few as 4.4% of mutations may be shared across sites^(14)^. Consequently, NDT based solely on neoantigens identified in the primary tumor may be ineffective for patients with metastatic disease. In contrast, targeting ubiquitous neoantigens–shared across all tumor sites–is an attractive approach, provided antigen presentation remains intact and systemic immune competence is preserved^(15, 16)^. Several clinical trials have demonstrated responses to NDT in metastatic sites, even for tumors with low tumor mutational burden (TMB)^(17, 18, 19, 20, 21, 22)^. In a clinical trial of adoptive T- cell therapy for metastatic colorectal cancer^(23)^, only T-cells targeting clonal neoantigens induced responses, supporting a focus on genetically conserved, widely shared neoantigen targets^(24)^.

We assessed here whether NDT might hold promise for patients with pancreatic neuroendocrine tumors (PNETs), where routine resection of primary and nodal metastases enables direct assessment of mutation sharing and immunogenicity. PNETs have low TMB, low tumor-infiltrating T-cells (TILs) and high recurrence rate, despite aggressive surgical and adjuvant treatments. Clinical trials investigating immune checkpoint blockade (ICB) showed limited success, underscoring the need for alternative strategies^(25, 26, 27)^.

Using comprehensive genomic and transcriptomic profiling of 14 tumor samples (primary and 13 synchronous metastases) from a treatment-naïve metastatic PNET, we assessed mutational status and presence of shared neoantigens. We identified ubiquitous neoantigen-reactive T-cells in peripheral blood and tracked associated clonotypes within multiple metastatic sites. Our findings provide direct evidence for the presence and expansion of ubiquitous neoantigen-reactive T-cells in a TMB-low, TIL-low tumor, highlighting the potential of NDT for metastatic PNETs and informing neoantigen selection strategies for solid tumors more broadly.

## METHODS

### Patient details

A female was diagnosed (Jun-2016) with non-functional, metastatic grade II PNET: Cytokeratin 7- negative (CK7^-)^, CK20^+^, CD56^+^, chromogranin^+^ and synaptophysin^+^, with 15-25% Ki67. At surgical resection (Aug-2016), prior to treatment, fresh and formalin-fixed, paraffin-embedded (FFPE) tissue were collected from 12 and 14 sites, respectively (Fig.1a, Supplemental Table S1). Primary tumor (site 1) comprised normal pancreatic tissue with a well-circumscribed nodule of neuroendocrine tumor. Nodal metastases (sites 2-14) showed extensive replacement by neuroendocrine tumor. Initial systemic therapy (Oct-2016 to Feb-2017; 5-Fluorouracil/Carboplatin/Streptozocin) achieved partial response. During surveillance, a leukapheresis product was collected (Sep-2018). Four cycles of peptide receptor radionuclide therapy (Sep-2018 to Jun-2019) achieved complete response. Upon disease progression (Sep-2020), FOLFOX (fluorouracil, leucovorin calcium and oxaliplatin) was administered (Apr to Jul-2022). Subsequent treatments included Everolimus (Mar to May-2023) and FOLFOX rechallenge (May to Jul-2023), until the patient’s death (May-2024).

**Figure 1.**
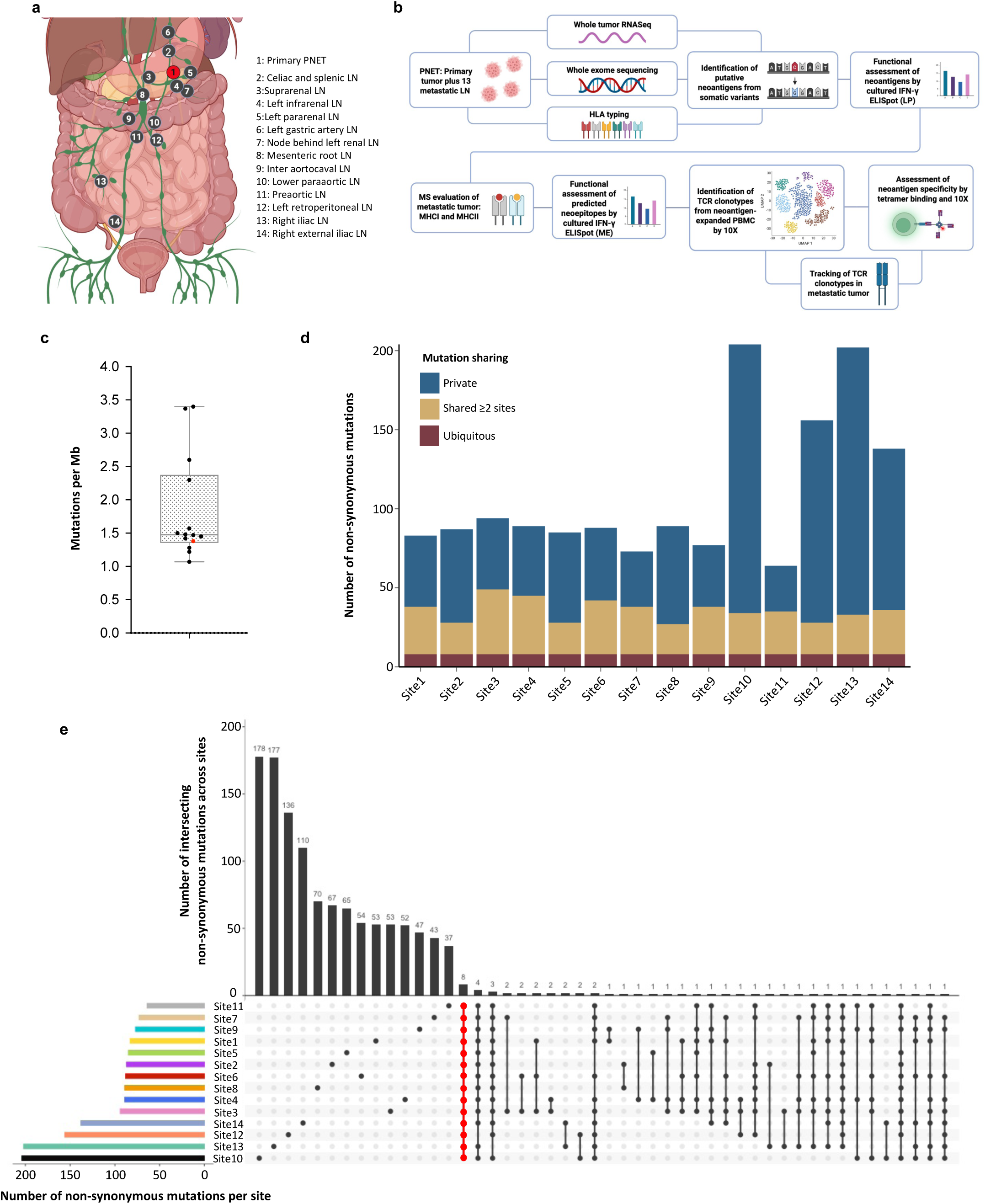
Mutational landscape of a patient with metastatic pancreatic neuroendocrine tumor (PNET) |. **a)** Schematic showing the location of the primary PNET (1, red circle) and 13 metastatic nodal sites (2-14, black circles). **b)** Multi-omic workflow used to identify neoantigens and associated reactive T-cell clonotypes, including WES, RNA/TCRseq, single-cell transcriptomics, immunopeptidomics and functional validation. **c)** Box plot showing the mutational burden of each tumor site: primary PNET (red circle) and 13 metastatic nodal sites (black circles). **d)** Bar chart showing the level of sharing of non-synonymous mutations across tumor sites: ubiquitous mutation (present in all sites, dark red bars), mutations shared by two or more sites (gold bars) and private (present in one single site, blue bars) are displayed. **e)** Detailed mutation sharing across tumor sites: upset plot (center) showing the sharing of non-synonymous mutations across tumor sites, vertical bars (top) indicate the number of private or shared mutations, colored horizontal bars (left) indicate the total number of mutations per site. Only the first 50 intersections are shown.

### Immunohistochemistry

Immunohistochemistry (IHC) of FFPE tumor (n=12/14, Supplemental Table S1) used antibodies: CD8a (clone C8/144B; DAKO), CD4 (clone 4B12; DAKO), CD103 (clone EPR4166(2); Abcam), Ki67 (clone MIB-1; DAKO), synaptophysin (clone DAK-SYNAP; DAKO), Major Histocompatibility Complex Class I (MHC-I, clone EMR8-5; Abcam) and Class II (MHC-II, clone CR3/34; Abcam). Sections were scanned using Zeiss Axio Scan.Z1 Digital Slide Scanner and Zeiss ZEN software (v2.6) and quantified in QuPath (v0.5.0) from three independent regions of interest (1×10^6^ µm^2)^.

### NGS and mutational analysis

DNA and RNA were extracted from 10-20 cryosections (10 µm) of snap-frozen tumor with the Maxwell RSC tissue DNA and simplyRNA tissue kits using a Maxwell RSC instrument (Promega). FFPE DNA was extracted with the Maxwell RSC DNA FFPE kit, and germline DNA from whole blood with the Maxwell RSC Blood DNA kit. Nucleic acid was quantified using a Qubit 4 Fluorometer (Thermo Fisher Scientific). RNA quality was assessed on an Agilent 2100 Bioanalyzer (Agilent Technologies UK Ltd.). DNA and RNA libraries were prepared using the Human All Exome V6 SureSelect XT2 (Agilent) and TruSeq Stranded mRNA (Illumina) Library Prep Kits, then 100-bp paired-end sequenced on an Illumina NovaSeq 6000 platform (average coverage 50–100X). Data are deposited in the European Genome-Phenome Archive (EGA; EGAS00001006722).

High-quality RNA sequencing (RNAseq) reads were aligned to the human genome GRCh38/hg38 reference using *STAR* (v2.0.9), processed with *HTSeq-count* (v0.5.4) and normalized as transcripts per million (TPM). Mean mapped reads was 78.6% (range 63.7-89.3%); poorly expressed genes (read count <10) were eliminated. The immune profile of metastatic tumors was contextualized using publicly available data: PNET (n=33, GEO under GSE118014) and head and neck squamous cell carcinoma (HNSCC, n=36, ArrayExpress under E-MTAB-4546)^(28, 29)^.

Whole exome sequencing (WES) reads were aligned to GRCh38/hg38 using *BWA* (v0.7.15). Duplicates were marked with *Picard* (v1.141). Mean mapped reads was 125M (range 23M-332M); 184M for germline. Somatic variants (non-synonymous) were identified with *Varscan2* (v2.3), *Strelka* (v2.8.4) and *Mutect* (v4.1.7.0) and annotated using *VEP* (v97) and *SNPeff* (3.1m). Variant calling was restricted to missense, single-nucleotide variants (SNVs) and in-frame indels, only if tumor allele frequency was >2% and if variants were detected by ≥2 variant callers (Supplemental Table S2). One discordant annotation (mutated *SEC16A2*) was resolved using a fourth variant caller (*SuperFreq*, v1.2.5).

### Evaluation of autologous T-cell responses to predicted neoantigens

Pre-existing T-cell responses to predicted neoantigens were assessed using cultured IFN-γ enzyme-linked immunosorbent spot assay (ELISpot) assay. Reactivity was first tested against mutation-derived long peptide (LP) pairs–20mers containing the altered amino-acid at position 6 (LP6) or 15 (LP15)–with corresponding wild-type (wt) peptides as controls (Supplemental Table S3).

Following four digit Human leukocyte antigen (HLA) typing (Supplemental Table S4) at NHS-BT H&I (http://hospital.blood.co.uk/diagnostic-services/hi/), mutation-derived minimal epitopes (ME) were predicted for mutated *NPTX2* (*mNPTX2*) and *SEC61A2* (*mSEC61A2)* using pVACtools (v.1.5.10). For MHC class I, for 8-11mer peptides predicted to bind HLA-A, B and C allotypes were selected; for MHC class II 15mer peptides predicted to bind to HLA-DRB1 were selected. Binding affinity prediction were performed using MHCflurry, MHCnuggetsI, NetMHC, NetMHCcons, NetMHCpan, PickPocket, SMM, and SMMPMBEC for HLA class I and MHCnuggetsII, NNalign, NetMHCIIpan, and SMMalign for HLA class II. For each algorithm, only ME with Inhibitory concentration 50 (IC50) <500 nM were retained. A median IC50 across all algorithms was then calculated for each epitope. mNPTX2 and mSEC61A2-derived ME with the lowest median IC50 were selected for T-cell stimulation (Supplemental Table S4). Peptides were synthesized to >95% (Peptide Protein Research Ltd.) and reconstituted in 100% DMSO.

For both LP and ME stimulations, peripheral blood mononuclear cells (PBMCs, 2×10^6^/well) from leukapheresis were stimulated with peptide (5 µg/mL) and recombinant Interleukin 2 (IL-2, 20 IU/mL; R&D Systems Europe Ltd.) in 24-well plates at 37°C, 5% CO_2_. Media containing IL-2 (20 IU/mL) was refreshed on days 4, 6, 8 and 11; cells were harvested on day 13. Expanded cells were either cryopreserved for single-cell RNA and T-cell receptor sequencing (scRNA/TCRseq, 10x Genomics) or underwent IFN-γ ELISpot. For ELISpot, 1×10^5^ cells/well were re-stimulated with peptide (5 µg/mL) in triplicate for 22 h; phytohemagglutinin (PHA; Sigma-Aldrich Company Ltd.) and CEFT (JPT Peptide Technologies GmbH) served as positive controls. Spot-forming cells (SFC) were counted using the AID ELISpot plate reader (AID Autoimmun Diagnostika GmbH) and response was classified using the runDFR(x2) tool (https://rundfr.fredhutch.org).

### NPTX2-specific tetramer staining

PBMC were thawed, rested for 24 h and stimulated in 24-well plates (2×10^6^/well) with mNPTX2.ME.HLA-I (FEWGNNPIEL, 10µg/mL) with recombinant IL-7 (10 ng/mL) and IL-15 (5 ng/mL; both Gibco) at 37°C, 5% CO_2_. Media was supplemented with IL-7 (10 ng/mL), IL-15 (5ng/mL) and IL-2 (20 IU/mL) on day 3, and IL-7 and IL-15 alone on days 6, 8, 9 and 10. Cells were harvested on day 11 and CD8^+^ T-cells were enriched using a CD8^+^ isolation kit (Miltenyi Biotech). Peptide-expanded, CD8-enriched cells were incubated with FcR blocking reagent (Miltenyi Biotech, 10 min, 4°C), followed by PE-labelled HLA-B*40:02-restricted MHC-I tetramer (FEWGNNPIEL, 6 µg/mL; Creative Biolabs, lot CB2308Y06, 30 min, room temperature). Additional staining (30 min, 4°C) was performed with antibodies (BioLegend): CD3-PE-Cy7 (clone UCHT1), CD4-FITC (clone RPA-T4), CD8-BV510 (clone SK1), CD69-BV421 (clone FN50), CD137-PE-Texas Red (clone 4B4-1) plus Fixable Viability Dye eFluor780 (Invitrogen).

Tetramer-positive cells were sorted into FACS buffer with 50% fetal bovine serum (FBS) and 0.2 U/mL RNase inhibitor on a BD FACSAria III cell sorter and analyzed in FlowJo (v10.8.1) before scTCRseq (10x Genomics). Controls included ex-vivo PBMCs and peptide-expanded or unstimulated PBMCs from a non-HLA matched healthy donor.

### Bulk TCR**β** repertoire analysis

Bulk TCRβ sequencing was performed on RNA from metastases (n=8/13). Libraries (TrueSeq Nano DNA, Illumina) were pooled and sequenced on a MiSeq (150-bp paired-end). Reads were processed with *MIGEC* (v1.2.9), and downstream analyses performed in R (v3.6.3) using *powerTCR* (v1.22.0) and *immunarch* (v0.9.1). TCRβ reconstruction succeeded for all samples (92% unique molecular identifiers (UMIs) mapped).

A clonotype was defined by a unique Vβ CDR3 amino-acid sequence. Clone size distributions were modelled with the Gamma–GPD spliced threshold model using *powerTCR*^(30)^ to define expansion thresholds. Highly expanded clonotypes were those exceeding the 95^th^ percentile of expanded clones. Diversity (Chao1 estimator) and clonotype overlap (Morisita–Horn index) were computed using *immunarch*.

### 10x Single-cell RNA/TCR analysis

#### Cell sorting

For mNPTX2.LP/ME-expanded PBMCs, thawed cells were rested for 2 h, blocked with FcR (Miltenyi, 10 min, 4°C), stained with TotalSeq C hashtags (BioLegend; 15 min, 4°C) and DAPI, and up to 60,000 live cells were sorted on a BD FACS ARIA II into phosphate buffer saline (PBS)/FBS (1:1) with RNase inhibitor (1:100; Takara). mNPTX2-tetramer T-cells were sorted similarly. Sorted cells were diluted with ice-cold PBS to 1.4 mL, centrifuged (600 g, 4 °C, 10 min), resuspended in 25 μL 0.04% BSA/PBS, and 33 μL was used for 10x Genomics processing following the manufacturer’s instructions.

#### 10x Genomic library preparation and quality control

scRNAseq libraries were prepared using the 10x Genomics 5′ TAG v2.0 chemistry with immune-profiling and cell-surface protein capture, following manufacturer protocols. cDNA and library amplifications used 13 cycles; V(D)J and ADT libraries used 9 and 8 cycles, respectively. Libraries were pooled equimolarly and sequenced on Illumina NovaSeq 6000 (read1/read2 = 101 cycles, i7 index = 8 cycles).

Reads were demultiplexed with *bcl2fastq* (v2.20.0.422) and aligned to GRCh38 (v3.0.0) using C*ell Ranger* (v7.1.0). Doublets were removed using hashtag demultiplexing and Scrublet (v0.2.3; score >0.3). Low-quality droplets (<200 or >4,500 genes, <400 or >20,000 UMIs, >10% mitochondrial UMIs) were excluded. Library quality control metrics are in Supplemental Table S5.

#### Differential gene expression analysis and cluster annotation

Genes expressed in ≥0.1% of cells were retained. Data were normalized and integrated in *Seurat* (v3.1.5)^(31)^, selecting variable genes (mean UMI count >0.01, top 25% variance) and the first 20 principal components for clustering using *FindNeighbors* and *FindClusters* functions. Clusters enriched in heat-shock^(32, 33)^ and B-cell genes (CD79A, immunoglobulins) were excluded. After testing multiple resolutions, 0.15 was chosen for downstream analyses.

Differential expression was computed with MAST (v1.10.0); genes with adjusted p-value < 0.05 and log (fold change) >0.35 were considered significant. Signature module scores were calculated as previously described^(34)^, averaging expression per cluster using gene lists^(35, 36, 37, 38)^ (Supplemental Table S6).

#### Single-cell TCR repertoire analysis

Single-cell V(D)J-enriched libraries were processed with the *Cell Ranger* vdj pipeline (v3.1.0, GRCh38 reference). Clonotypes were defined by unique α, β, or paired αβ CDR3 amino-acid sequences. Clonotypes detected in ≥2 cells were considered expanded, assessed globally and per cluster. Clone size per cluster was visualized on uniform manifold approximation and projection (UMAP) and diversity (Chao1 estimator) calculated with *immunarch*.

#### Cell type identification

Antibody-derived tag (ADT) matrices for CD4 and CD8B (TotalSeq-C, BioLegend; Supplemental Table S7) were CLR-normalized in *Seurat* (v4.3.0). Expression distributions were modeled using a three-component Gaussian Mixture Model (negative, low, high) in *Mclust* (v6.1.1). Thresholds between consecutive components defined CD4, CD8, double-negative (DN), or double-positive (DP) subsets.

For 25 clonotypes shared between CD4act and CD8act clusters, ADT and gene expression (GEX) data were integrated to refine classification: cells expressing CD8B (GEX and/or ADT) only were labeled CD8, CD4 (GEX and/or ADT) only as CD4, both as DP (GEX only), and neither as DN (Supplemental Table S8). After excluding DN/DP cells, 22/25 shared clonotypes were exclusively CD4 or CD8; the remaining three, with <0.5% minor populations, were assigned based on the dominant phenotype.

### Clonotype overlap analysis

Clonotype overlap was assessed between scTCRseq datasets (mNPTX2 LP/ME-expanded and tetramer T-cells) and bulk TCRβ repertoires from metastatic sites. For scTCRseq, only expanded paired clonotypes were included, and overlap was defined as an identical amino-acid CDR3 sequence for both chains. When comparing single-cell with bulk data, only Vβ CDR3s were used. If several single-cell clonotypes overlapped with a single bulk clonotype (e.g., same β, different α), all were counted. Conversely, when one single-cell clonotype overlapped with multiple bulk TCRβ clonotypes (dual β chains), only the bulk clonotype detected in the most metastatic sites was retained.

### Mass spectrometry

Snap-frozen tumor samples were thawed, weighed, and homogenized in 4 mL lysis buffer (0.02 M Tris, 0.5% IGEPAL, 0.25% sodium deoxycholate, 0.15 mM NaCl, 1 mM EDTA, 0.2 mM iodoacetamide, protease inhibitors)^(39)^. Homogenates were clarified (2,000 g, 10 min and 13,500 g, 60 min, 4 °C). MHC-I complexes were immunoprecipitated with 2 mg anti-MHC-I mouse monoclonal antibody (W6/32) covalently linked to Protein A–Sepharose (Repligen) for 2 h, 4 °C, washed in isotonic (0.15 M NaCl) and hypertonic (0.4 M NaCl) Tris-buffered saline, eluted with 10% acetic acid and dried under vacuum. MHC-II peptides were captured from the MHC-I–depleted lysate using anti-MHC-II mouse monoclonal antibody (IVA12)-conjugated beads and eluted identically.

Eluates were reconstituted in buffer (1% acetonitrile, 0.1% trifluoroacetic acid), loaded onto pre-wet HLB columns (Waters) and washed with 5 column volumes of 0.1% TFA. Immunopeptides were separated from MHC-I/β2m or MHC-II heavy chain using 12 sequential step elution (0.5 mL) from 0- 30% acetonitrile in 0.1% TFA. Odd and even fractions were pooled and dried under vacuum. Immunopeptide fractions were reconstituted in 0.1% formic acid, split into two and separated by liquid chromatography-tandem mass spectrometry (LC-MS/MS) on an Ultimate 3000 RSLC nano system (Thermo Fisher Scientific) using a PepMap C18 EASYSpray LC column (2 μm particle size, 75 μm x 75 cm; Thermo Fisher Scientific) in buffer A (0.1% formic acid) and coupled on-line to an Orbitrap Fusion Tribrid Mass Spectrometer (Thermo Fisher Scientific) with a nano-electrospray ion source. Peptides were eluted with a linear gradient of 3-30% buffer B (acetonitrile and 0.1% formic acid) at a flow rate of 300 nL/min over 110 min. Immunopeptide fractions were first analyzed using an untargeted data-dependent approach, followed by targeted analysis using prioritized peptide lists (Supplemental Tables S9–S10)

#### Untargeted immunopeptide analysis

Untargeted LC-MS/MS acquisition was performed in Top Speed data-dependent mode with one full MS scan every 3 s, followed by higher energy collision-induced dissociation MS/MS. MS spectra were acquired in the Orbitrap at 120,000 resolution (300 m/z), with an automatic gain control ion target value of 4.0×10^5^ with a maximum of 100 ms. MS spectra were acquired at 30,000 resolution (100 m/z). HCD energies were 28 for charge 2–4 and 32 for singly charged peptides. Fragment ions were analyzed in the Orbitrap, and previously fragmented precursors dynamically excluded for 30 s.

#### Targeted immunopeptide analysis

Targeted LC-MS/MS was performed on the second peptide fraction using a prioritized inclusion list (charge states 1-4, intensity threshold 20,000; Supplemental Tables S9–S10), using identical fragmentation parameters to the untargeted run. A secondary scan at lower priority captured additional peptides with charge states 2-4.

### Mass spectrum data analysis

Raw spectra were processed in *Peaks Studio* (v10.0 build 20190129) to generate reduced charge state and deisotoped precursor and product ion lists. Searches were performed against UniProt (20,350 entries, 2020-04-07) supplemented with sample-specific mutanomes (1,000-5,000 sequences) and contaminants, using unspecific digestion^(39)^. Parent and fragment mass error tolerances were 5 ppm and 0.03 Da, respectively. Variable modifications included N-terminal acetylation (42.01 Da), methionine oxidation (15.99 Da), and cysteine carbamidomethylation (57.02 Da). Carbamidomethylation was set as variable due to low iodoacetamide concentration (0.2 mM) used. Up to three variable modifications per peptide were allowed. False discovery rate (FDR) was 1% using decoy-fusion searches. Downstream analyses and visualization were performed in R. Data are available via PRIDE (PXD037449; DOI 10.6019/PXD037449)^(40)^.

### Data visualization

Data visualization was performed in R using *ggplot2* (v3.5.2) for bar, scatter, box, and heat maps; *Seurat* (v5.2.1) for UMAPs and dot plots; *immunarch* (v0.9.1) for TCR diversity plots; and *UpSetR* (v1.4.0) for clonal overlap. Illustrative schematics were created with BioRender.com.

## RESULTS

Tissue from the primary PNET (site 1), from 13 metastatic lymph nodes (sites 2-14; Fig.1a) and PBMCs, were analyzed using a multi-omics workflow to identify neoantigens and neoantigen-reactive T-cell clonotypes (Fig.1b; Supplemental Table S1).

### Tumor characterization

Histopathology confirmed a well-differentiated grade II PNET with extensive nodal metastases (Supplemental Fig.S1a). Tumor cells strongly expressed synaptophysin and MHC-I, while MHC-II (median 16.8%) was detected on endothelial, histiocytic, lymphoid, and some tumor cells (Supplemental Table S11). T-cell infiltrate was low, with a median 4.0% and 0.9% of CD4 and CD8 T-cells, respectively, and cells were mostly located in the stroma between tumor cell nests. CD103^+^ T-cells were rare (0.2%). All metastases showed low TILs with similar distribution and no evidence of immune exclusion.

Using whole-tumor RNAseq and previously published data for contextualization^(28, 29)^, *CD4* and *CD8* gene expression was low (Supplemental Fig.S1b), corroborating immunohistochemistry results. CIBERSORT showed median CD4^+^ and CD8^+^ T-cell proportions of 38.5% and 4.2%, respectively (Supplemental Fig.S1c). MHC-I-related genes (e.g. *HLA-A/B/C*, *B2M*, *PDIA3*) were abundantly expressed while MHC-II-related genes (e.g. *CIITA*, *HLA-DOB*, *IFI30)* were expressed at lower levels (Supplemental Fig.S1d), consistent with immunohistochemistry.

### Tumor mutational landscape

WES of the primary tumor and metastases identified 1,195 unique nonsynonymous mutations. The median number per site was 88 (range 64-204), corresponding to a low TMB (1.48 Mut/Mb, range 1.07-3.4; sequencing target size 60Mb, Fig.1c) consistent with previous reports^(41)^. Eight missense mutations were ubiquitous across all 14 sites and eight were shared by 13/14 sites (Fig.1d-e). Ubiquitous mutations represented 3.9-12.5% of total mutations per site, including 9.6% in the primary tumor (Fig.1d; Supplemental Table S12). Mutations shared by ≥2 sites accounted for 12.4-43.6% per site. No alterations were found in *ATRX*, *DAXX*, or *MEN1*, indicating an A–D–M wild-type phenotype^(28)^.

### Evaluation of T-cell reactivity to putative neoantigens–long peptides

We next evaluated the immunogenicity of neoantigens derived from ubiquitous mutations using cultured IFN-γ ELISpot assay. Of eight ubiquitous mutations, seven were tested (Supplemental Table S3); *MUC16* was excluded due to low expression (Supplemental Table S13). Autologous T-cell responses were observed exclusively to long peptides derived from *mNPTX2* and *mSEC61A2* (Fig.2a); reactivity was specific to LP6 peptide for mNPTX2, while both LP6 and LP15 generated responses for mSEC61A2. The remaining candidates showed no reactivity (*SPPL3*, *NOTCH3*), equivalent reactivity to mutant and wt (*LRRC8D*, *CEP295*) or reactivity to the wt alone (*ULK1*), indicating that only mNPTX2 and mSEC61A2 generated neoantigen-specific responses.

**Figure 2.**
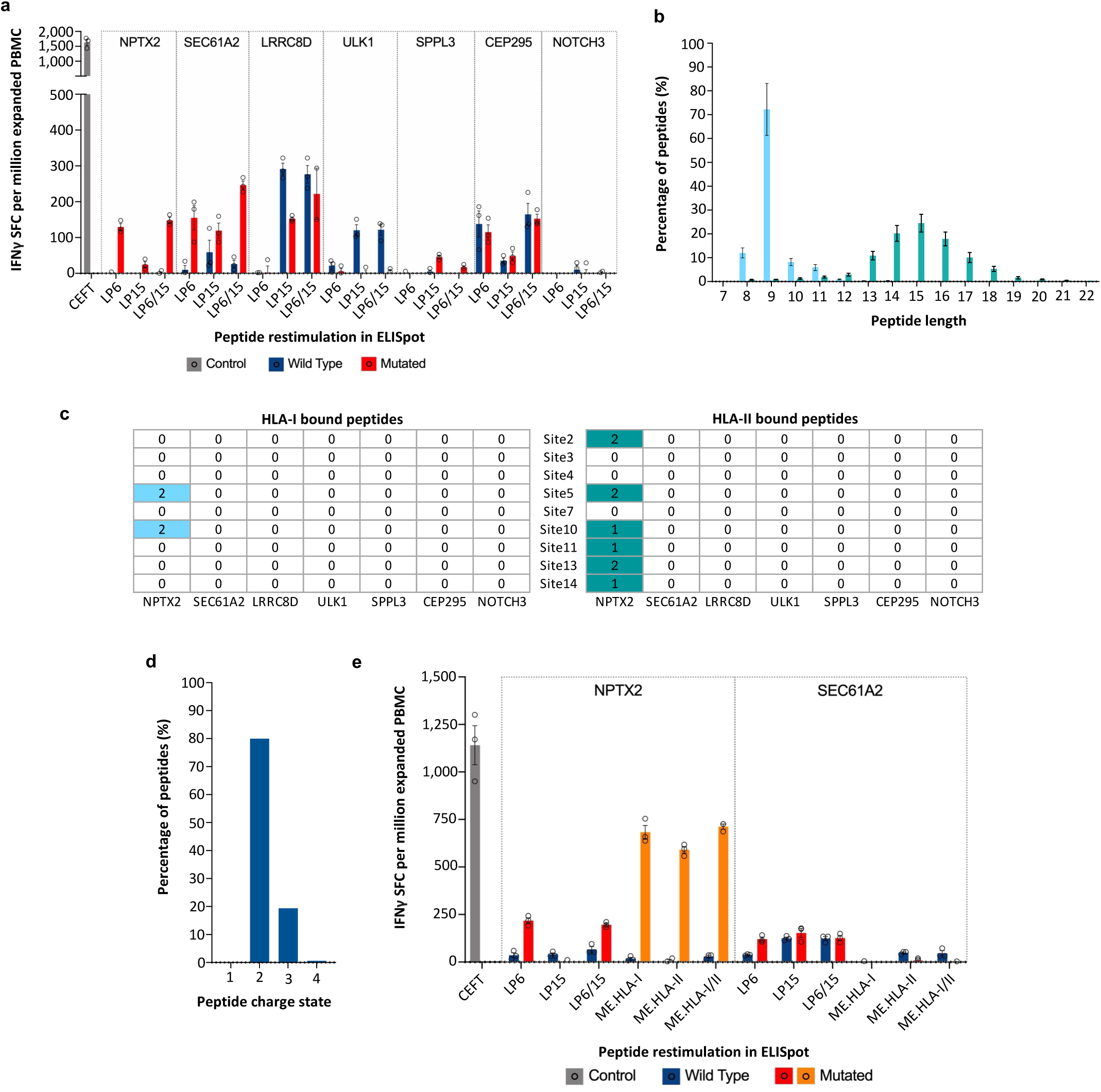
Identification of potential neoantigens |. **a)** Bar plot showing cultured IFN-γ ELISpot response in spot forming cells (SFC) per million expanded PBMC. The color of the bar represents the peptide used: synthetic mutated long peptides (red bars); LP6 means the peptide is associated with altered amino-acid in position 6, LP15 means the peptide is associated with altered amino-acid in position 15, wild type peptides (blue bars). Viral peptide pool (CEFT, grey bar) used as a positive control. Each dot represents an individual experiment performed in triplicate. Data expressed as mean ± standard error of the mean (SEM) of triplicate. **b)** Bar plot showing the proportion of peptides (y axis) for each peptide length (x axis) for HLA-I (n=18,340, blue bars) and HLA-II (n=2,082, teal bars) eluted from metastatic tumor (n=9 sites). **c)** Tables showing the number of HLA-I (blue) and HLA-II (teal) bound peptides captured on mass spectrometry for each candidate neoantigen across the 9 sites analyzed. **d)** Bar plot showing the proportion of peptides for each charge state. **e)** Bar plot showing the cultured IFN-γ ELISpot response in SFC per million of expanded peripheral PBMCs. The color of the bar represents the peptide used: synthetic mutated minimal epitope peptides (orange bars), synthetic mutated long peptides (red bars), wild type peptides (blue bars). Viral peptide pool (CEFT, grey bar) was used as a positive control. Each dot represents an individual experiment performed in triplicate. Data expressed as mean ± SEM of triplicate.

### MHC peptide elution from metastatic tumor

To define tumor immunopeptidomes, peptides were eluted from MHC molecules of nine metastatic sites and analyzed by LC-MS/MS. We identified 18,340 HLA-I and 2,082 HLA-II peptides (1% FDR) with characteristic length distributions (Fig.2b). Peptides derived from the seven predicted ubiquitous neoantigens were not observed. However, wtNPTX2-derived peptides were captured in two and six metastatic sites for HLA-I (DFREVLQQR, GELERQLL) and HLA-II (INDKVAQLPLFVSDG, INDKVAQLPLFVSDGK), respectively (Fig.2c). mNPTX2-derived peptides were not captured, likely because they were predicted to be singly charged (Supplemental Fig.S2a), an ionization state typically underrepresented in LC-MS/MS datasets^(42, 43)^. In agreement, no singly charged peptides were detected in our data (Fig.2d) and the observed wtNPTX2 peptides exhibited optimal charge states (2 and 3) (Supplemental Fig.S2b). Nonetheless, the frequent detection of wtNPTX2-derived peptides suggested possible presentation of the mutant counterpart.

### Evaluation of T-cell reactivity to putative neoantigens–minimal epitope

The mNPTX2- and mSEC61A2-derived ME with the highest median binding affinity, along with their wt counterparts, were synthesized for IFN-γ ELISpot assay. Both mNPTX2.ME.HLA-I and mNPTX2.ME.HLA-II elicited a strong response, 2-3-fold higher than those to the LP, while no response was observed to the equivalent wt peptides. In contrast, mSEC61A2 failed to elicit significant responses with either ME or LP (Fig.2e).

### TCR analysis in metastatic sites

To assess whether mNPTX2-reactive T-cell clones were present in tumor tissue, we first profiled the TCRβ repertoire of eight metastatic sites by bulk TCR sequencing (Supplemental Table S14). This analysis provided a reference map for subsequent identification of reactive clonotypes. Per site, the median number of clonotypes was 272 (range 221-442) and UMI count was 4,445 (range 2,243- 7,950) (Supplemental Fig.S3a). Diversity (Chao1 estimator) and clonal distributions were comparable across sites (Fig.3a–b). Using the Gamma–GPD spliced threshold model, expanded and highly expanded clonotypes were defined (Fig.3c; Supplemental Fig.S3b); over half of all clonotypes were expanded per site (51–82.3%; Fig.3d). TMB and inferred total, CD8 and CD4 T-cell abundance did not correlate with clonotype richness, expansion or diversity (data not shown). While most clonotypes were site-specific (Fig.3e-g), a subset was shared across metastases (e.g., sites 5 and 11, Morisita– Horn index = 0.19, Fig.3g), suggesting potential recognition of antigens shared across tumor sites.

**Figure 3.**
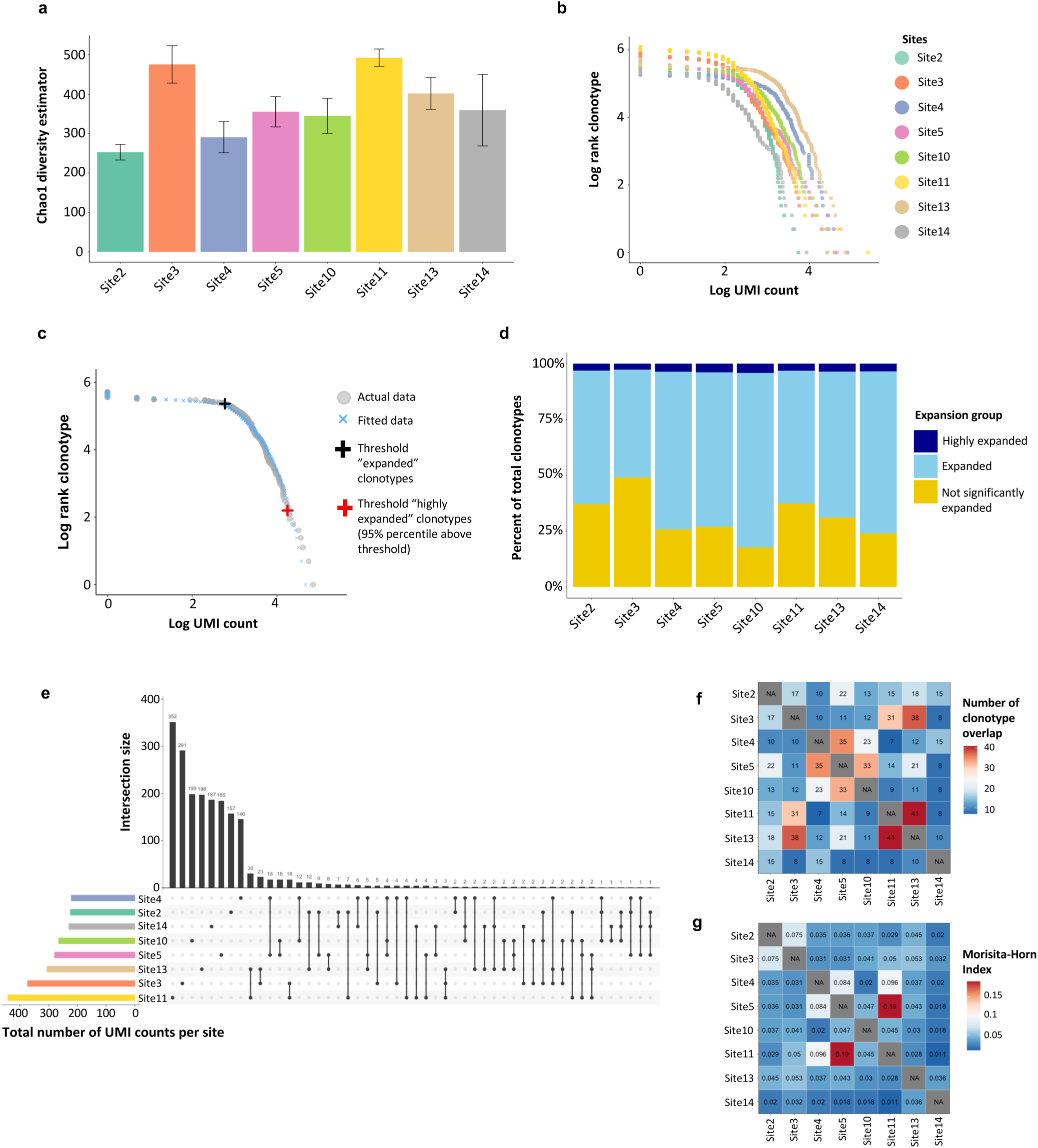
TCR evaluation of nodal metastases |. **a)** Bar plot showing the estimated TCRβ diversity (Chao1 estimator) of clonotypes across metastatic sites (n=8). Error bars represent the standard deviation (SD) of the Chao1 estimates, which were calculated using a bootstrapping procedure to account for sampling variability. **b)** Plot showing TCRβ UMI count distribution across metastatic sites (n=8). **c)** Plot showing TCRβ UMI count distribution for one representative metastatic site: thresholds for “expanded” (black cross) and “highly expanded” (red cross) clonotypes are shown. **d)** Bar plot showing the proportion of “highly expanded” (dark blue bars), “expanded” (light blue bars) and “not significantly expanded” (yellow bars) clonotypes for each metastatic site. **e)** TCRβ sharing across metastatic sites: upset plot (center) showing clonotype sharing across tumor sites, black vertical bars (top) indicate the number of private or shared clonotypes across sites, colored horizontal bars (left) indicate the total number of clonotypes per site. Only the first 50 intersections are shown. **f)** Heatmap showing TCRβ clonotype sharing; values correspond to the number of shared clonotypes identified between two sites (color scale). **g)** Heatmap of similarity index (Morisita-Horn index) of TCRβ clonotypes across metastatic sites; values correspond to the Morisita Horn index identified between two sites (color scale).

### Single-cell RNAseq and TCR analysis of mNPTX2-expanded PBMCs

To identify mNPTX2-reactive clonotypes, autologous PBMCs were stimulated with mNPTX2- derived LP (LP6/LP15) and ME (HLA-I/II) pools in separate cultures for 13 days, then sorted and analyzed by scRNA/TCRseq (Fig.4a–b). After quality control steps to remove low-quality cells, doublets, and B-cells, 35,178 cells were retained (Supplemental Fig.S4a-f, Supplemental Table S5, S15; 64% had productive TCRs yielding 4,289 clonotypes, of which 2,701 had paired Vα/Vβ information.

**Figure 4.**
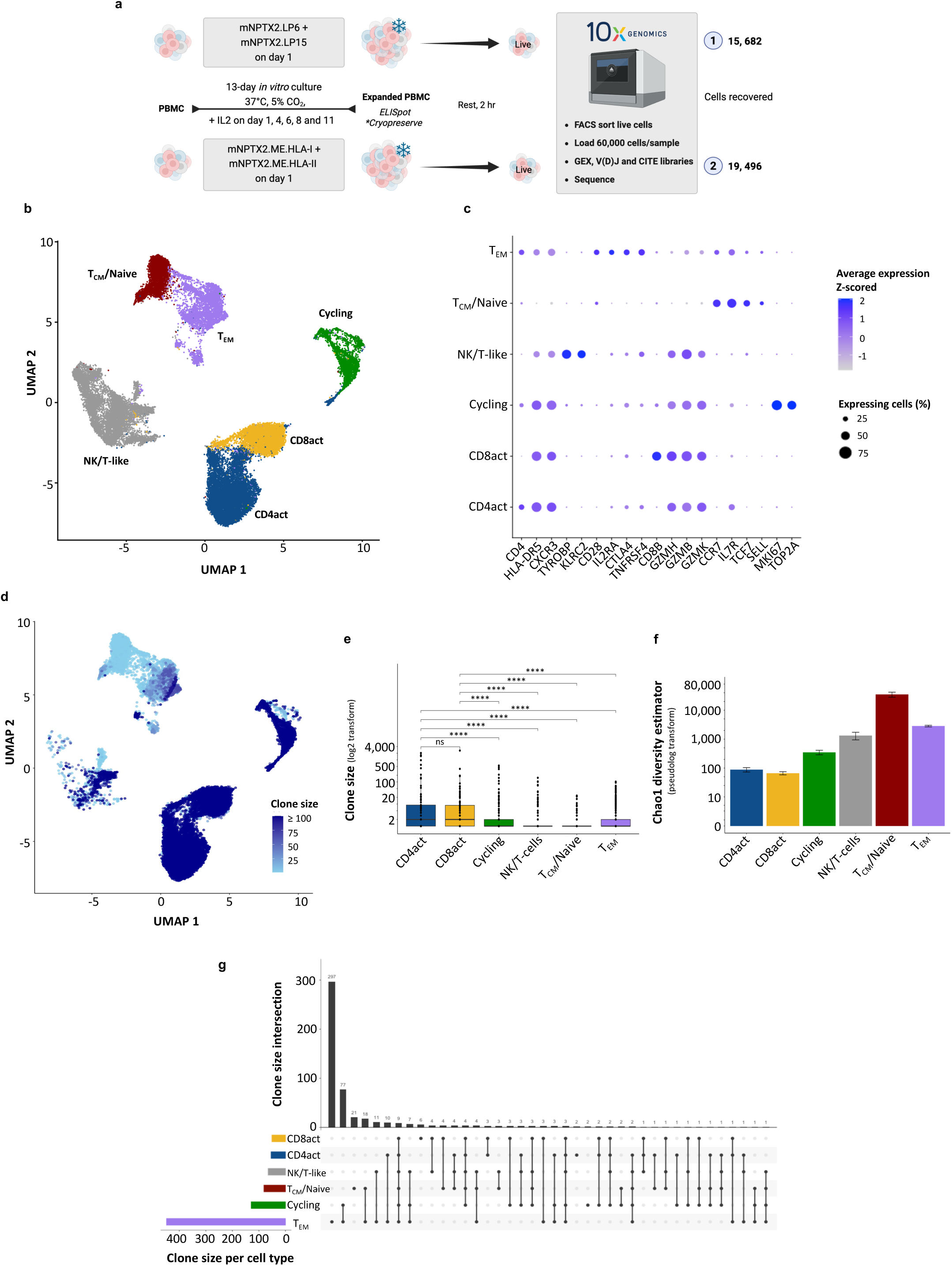
Single-cell transcriptomics of mNPTX2-expanded T-cells |. **a)** Schematic showing the experimental workflow used for sc RNA/TCRseq of mNPTX2-expanded autologous T-cells. **b)** UMAP of the single-cell transcriptome of mNPTX2-expanded T-cells showing 6 distinct clusters. **c)** Dot plot showing the Z-score-normalized mean transcript expression (color scale) and percentage (size scale) of expressing cells for significantly enriched relevant genes in each cluster. **d)** UMAP showing the distribution of TCR clone sizes (color scale). **e)** Box plot showing median clonotype size (log_2_ scale) across clusters. Statistical significance between clusters was assessed using the two-sided Wilcoxon test with p-values adjusted for multiple comparisons using the Benjamini-Hochberg method. **** indicates a p-value < 0.0001. **f)** Bar plots showing the estimated TCRβ diversity (Chao1 estimator) of clonotypes across different clusters. Error bars represent the SD of the Chao1 estimates, which were calculated using a bootstrapping procedure to account for sampling variability. **g)** Clonotype sharing across clusters: upset plot (center) showing clonotype sharing across clusters, vertical bars (top) indicate the number of private or shared clonotypes, colored horizontal bars (left) indicate the total number of clonotypes per cluster. Only the first 50 intersections are shown. Only expanded clonotypes (clone size ≥2) and with paired αβ TCR chains are shown.

Transcriptomic clustering revealed six subsets, annotated based on the most differentially expressed genes (Fig.4c; Supplemental Fig.S4g, Supplemental Table S16) and gene signature scoring (Supplemental Fig.S4h): activated CD8 (CD8act; *CD8, GZMH, GZMB*), activated CD4 (CD4act; *CD4, HLA-II genes, GZMB*), cycling (*MKI67, TOP2A*), NK/T-like (*TYROBP, KLRC2*), central memory/naïve (T_CM_/Naïve*; TCF7, CCR7, SELL*) and effector memory (T_EM_*; IL7R, CD28, CCR7*). The most expanded clonotypes mapped predominantly to CD8act and CD4act clusters, which displayed larger clone sizes and lower diversity (Chao1) than other clusters (Fig.4d–f), indicating antigen-driven proliferation.

Twenty-five clonotypes were shared across CD4/CD8 activated clusters (Fig.4g) but associated cells showed clear skewing toward one lineage based on ADT data (Supplemental Fig.S5a-c). Integration of both ADT and GEX data confirmed this: 18 CD8act/CD4act shared clonotypes were defined as CD8 and 7 as CD4 (Supplemental Fig.S5d, Supplemental Table S8).

UMAP of LP- and ME-stimulated T-cells were mostly identical (Supplemental Fig.S6a), indicating that both stimulation conditions elicited similar transcriptional programs. In addition, 4.3% of clonotypes were shared, which represented 73.5% of recovered cells, predominantly mapping to CD8act and CD4act clusters (Supplemental Fig.S6b). The median clone size of these shared clonotypes was significantly larger than clones unique to either dataset, with no size difference between LP and ME conditions (Supplemental Fig.S6c), further supporting that shared clones, and by extension the CD8act and CD4act clusters, were enriched for bona fide mNPTX2-reactive T-cells.

### Identification of mNPTX2-reactive CD8^+^ T-cell clonotypes

To validate that CD8act CD8^+^ clonotypes were mNPTX2-reactive, we isolated HLA-B*40:02- restricted mNPTX2.ME.HLA-I CD8^+^ T-cells using tetramer staining (Supplemental Fig.S7a-b). Upon scTCRseq of 16,000 sorted tetramer^+^ CD8^+^ T-cells, 3,783 cells and 1,619 unique paired clonotypes were recovered after quality control; 239 clonotypes were expanded (≥2 cells), accounting for 2,403 cells (Supplemental Fig.S7c).

Expanded tetramer-positive clonotypes were overlapped with those from mNPTX2.LP/ME cultures (Fig.5a). The majority of shared clonotypes mapped to the CD8act cluster, representing 95% of cells and 50% of clonotypes. Restricting analysis to ADT- and GEX-validated CD8^+^ clonotypes increased the overlap to 96.6% of cells and 57.6% of clonotypes (Supplemental Fig.S8a). Shared clonotypes had significantly larger median clone sizes than non-overlapping clonotypes, and overlap mostly comprised of highly expanded clones across both datasets (Supplemental Fig.S8b–c). This further supported antigen-driven expansion and enrichment of mNPTX2-reactive CD8 T-cells within the CD8act cluster.

**Figure 5.**
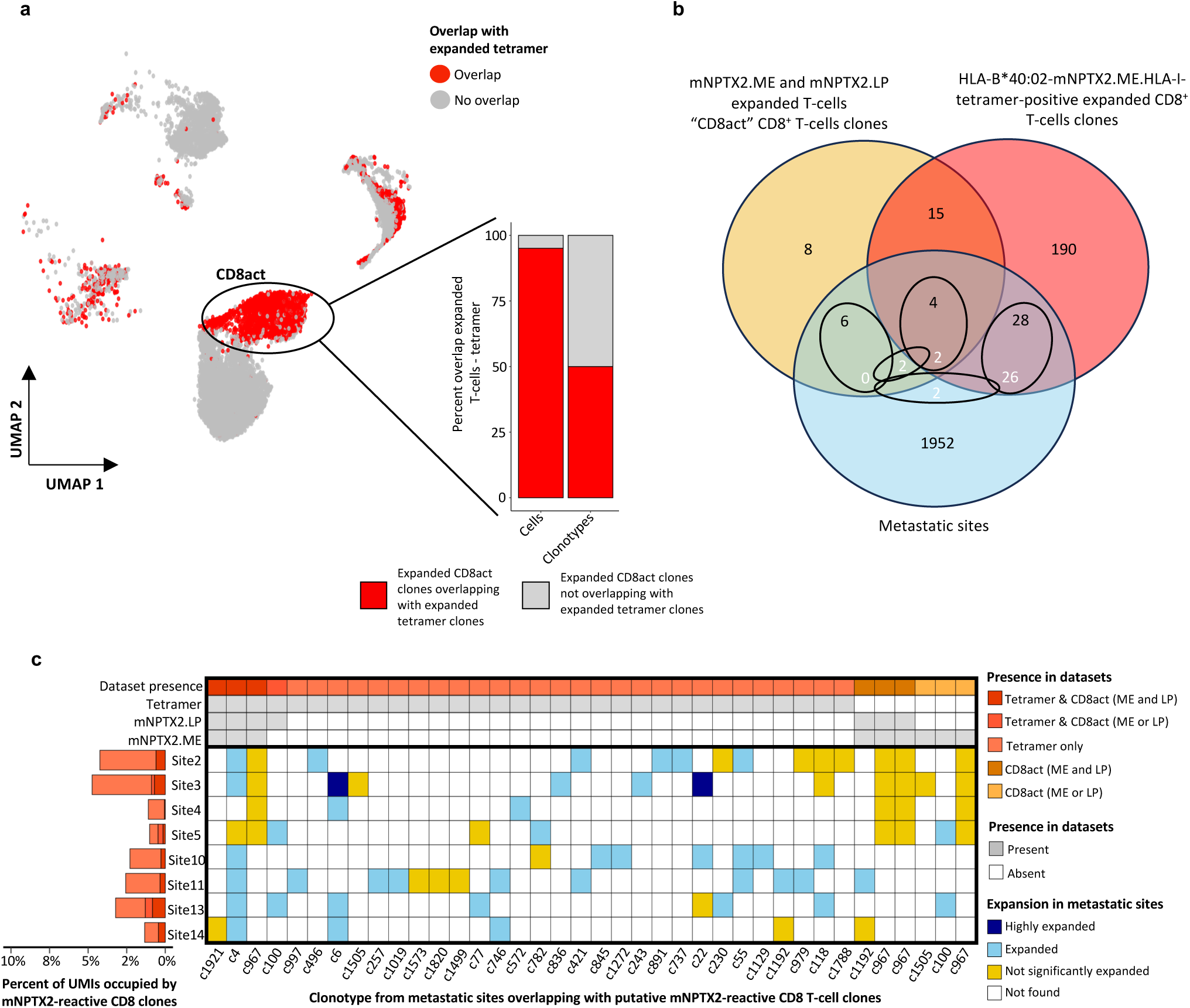
Identification of mNPTX2-reactive CD8^+^ T-cell clones |. **a)** UMAP of the single-cell transcriptome of mNPTX2-expanded T-cells showing the clonotype overlap (red circles) with tetramer-positive CD8^+^ T-cells clones. Only expanded clonotypes (clone size ≥2) are shown. The right bar plot shows the proportion of cells and clonotypes from the “CD8act” cluster that overlap with expanded tetramer-positive T-cells clones (red bars). **b)** Venn diagram showing clonotype overlap between: mNPTX2-expanded T-cells clonotypes (scTCRseq expanded, paired, CD8act cluster, CD8^+^ only), CD8^+^ tetramer^+^ T-cells clonotypes (scTCRseq, expanded, paired) and metastatic sites (bulk TCRseq, TCRβ only). In the inner circles, the upper values refer to the number of clones from scTCRseq overlapping with metastatic sites (n=38) and the lower (white) values refer to the clones from the metastatic sites. **c)** Summary of putative CD8^+^ mNPTX2-reactive clones across datasets. The top heatmap details the presence of each clone in the three scTCRseq datasets considered (LP6/LP15, ME.HLAI/II and tetramer^+)^. The bottom heatmap shows, for each clonotype, the level of expansion in each metastatic site (n=8). In both heatmaps, each row is a unique clone from scTCRseq and the clonotype ID at the bottom refers to clones in metastatic sites. The left bar plot shows the proportion of UMIs occupied by putative mNPTX2 reactive CD8 T-cell clonotypes in each metastatic site.

### Tracking of mutated mNPTX2-reactive clones in metastatic sites

To determine whether mNPTX2-reactive T-cells identified in peripheral blood were present in tumor tissue, we tracked all putative mNPTX2-reactive clonotypes–defined as expanded tetramer^+^ clonotypes and expanded CD8 clonotypes mapping to the CD8act cluster from the mNPTX2.LP/ME dataset–across bulk TCRβ repertoires from eight metastatic sites. We identified 38 putative mNPTX2-reactive CD8^+^ T-cell clonotypes across metastatic sites, including four shared between tetramer and mNPTX2.LP/ME datasets. In some instances, distinct scTCRseq clonotypes sharing a TCRβ chain overlapped with the same TCRβ clonotype in tumor tissue, yielding 32 unique TCRβ sequences (Fig.5b). For each clonotype, the number of metastatic sites containing putative mNPTX2-reactive T-cell clones ranged from 1 to 7, with most found to be expanded in tumor tissue and representing between 1.0-4.7% of TCR UMI counts per site (Fig.5c; Supplemental Table S17). Since MHC-II tetramers were unavailable, tracking of mNPTX2-reactive CD4 clonotypes was limited to bioinformatic analyses. After excluding misclassified CD8 clones, eight CD4 clonotypes, corresponding to seven unique TCRβ sequences, were identified across metastatic sites (Fig.6a), most of which were expanded (Fig.6b), representing 0.02–1.6% of TCR UMI counts per site. Overall, putative mNPTX2-reactive CD8^+^ and CD4^+^ T-cell clonotypes were detected in all metastatic sites analyzed, representing 1.5-5.1% of TCR UMI counts per site (Fig.6c).

**Figure 6.**
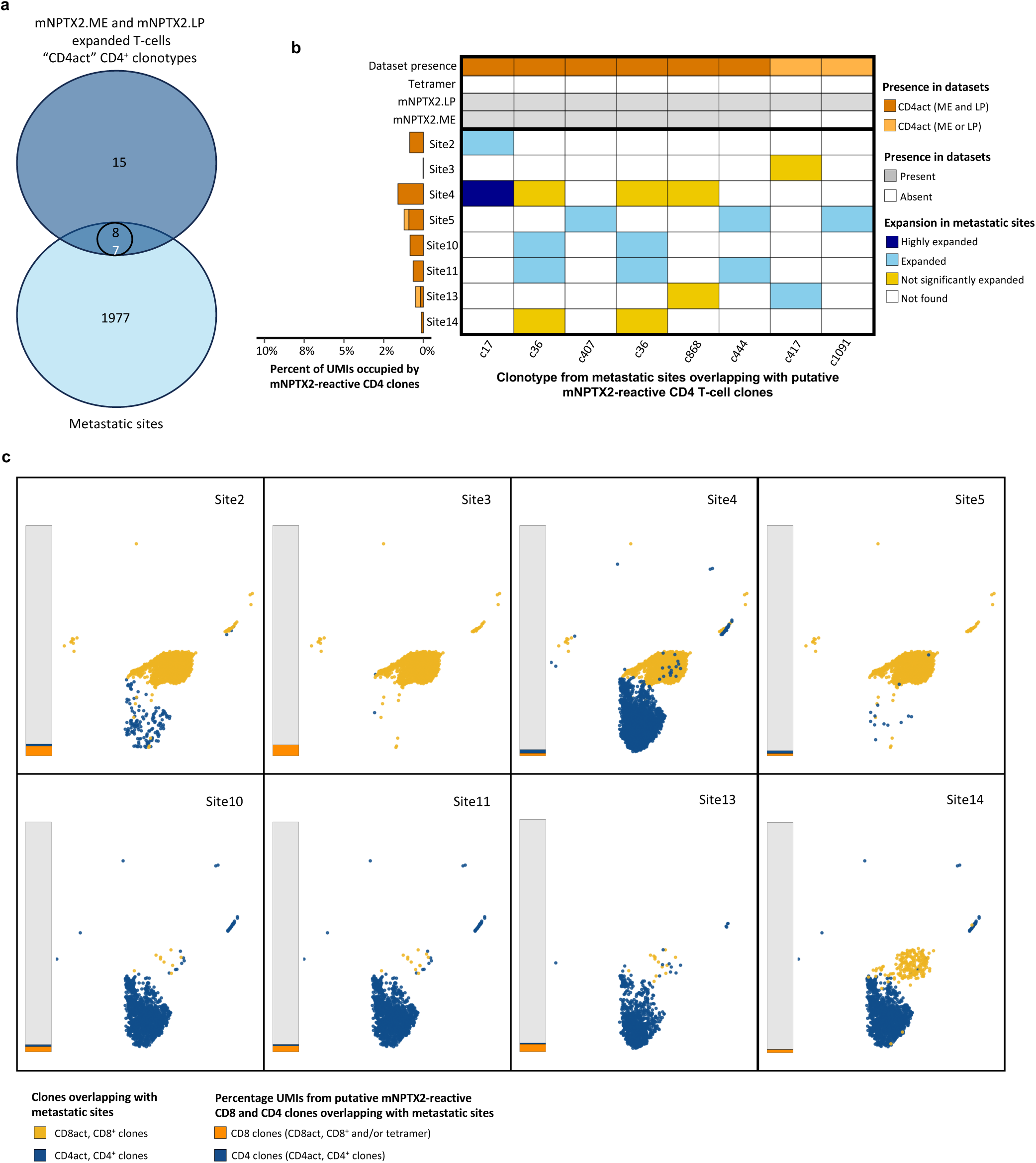
Identification of mNPTX2-reactive CD4^+^ T-cell clones |. **a)** Venn diagram showing clonotype overlap between mNPTX2-expanded T-cells clonotypes (scTCRseq, expanded, paired, CD4act cluster, CD4^+)^ and metastatic sites (bulk TCRseq, TCRβ only). In the inner circle, the upper value refers to the number of clones from scTCRseq overlapping with metastatic sites (n=8) and the lower (white) value refers to the clones from the metastatic sites. **b)** Summary of putative CD4^+^ mNPTX2-reactive clones across datasets. The top heatmap details the presence of each clone in the two scTCRseq datasets considered (LP6/LP15 and ME.HLA-I/II). The bottom heatmap shows, for each clonotype, the level of expansion in each metastatic site (n=8). In both heatmaps, each row is a unique clone from scTCRseq and the clonotype ID at the bottom refers to clones in metastatic sites. The left bar plot shows the proportion of UMIs occupied by putative mNPTX2 reactive CD4 T-cell clonotypes in each metastatic site. **e)** UMAP showing “CD8act” (yellow) and “CD4act” (blue) T-cell clones overlapping with each metastatic site (n=8). The left bar plot indicates the proportion of TCR UMIs occupied by putative mNPTX2-reactive CD8 (orange bars) and CD4 (blue bars) T-cells for each metastatic site.

## DISCUSSION

In a patient who presented with resectable multi-site metastatic, TMB low, TIL low, PNET, we detected eight ubiquitous mutations present in all tumor sites. Of these, one mutation-derived neoantigen elicited T-cell response in peripheral blood and pre-existing neoantigen-reactive T-cell clones could be tracked to all metastatic sites analyzed.

Of nearly 1,200 mutations detected, only eight were found to be ubiquitous. Although some mutations may have been missed, this low frequency of ubiquitous mutations was unexpected as recent evidence suggested that metastases are an early event in PNETs, with common shared mutations across metastatic sites^(44)^. Given the effect of treatment and time on immunoediting, mutation sharing was also expected to be higher in a treatment-naïve patient with synchronous nodal metastases^(11, 45)^ and perhaps the high number of tumor sites analyzed (n=14) contributed. Nonetheless, among eight ubiquitous mutations, one mutation-derived neoantigen was found to be immunogenic in peripheral blood and neoantigen-reactive clones could also be tracked across multiple metastatic sites, with evidence of expansion.

To identify neoantigen-reactive clones, we used a combination of scRNA/TCRseq after in-vitro expansion and scTCRseq of tetramer^+^ T-cells, enhancing detection sensitivity. We detected a total of 32 CD8 and 7 CD4 neoantigen-reactive TIL clones recovered from independent experiments. The observed polyclonal response to one neoantigen was consistent with previous data^(11, 46, 47, 48)^, most likely reflecting TCR promiscuity^(49)^. It has also been reported that T-cells might be primed by microbial antigens and subsequently react to neoantigens; this might add to the polyclonality of the observed T-cell response^(50, 51)^. Of note, we only considered clonotypes that were able to recirculate as we used PBMC to study the reactive repertoire. Therefore, we may have underestimated the true degree of polyclonality of mNPTX2-reactive T-cells.

Since the apheresis sample we used to probe circulating neoantigen-reactive T-cells was collected two years after tissue sampling, we evaluated peripheral neoantigen T-cell reactivities that have persisted over time. Our data therefore suggest that the mNPTX2 neoepitope has persisted. A lack of ELISpot response to the other six ubiquitous neoantigens may then reflect either lack of presentation, poor immunogenicity or neoantigen loss^(52)^.

Between metastatic sites, both overlapping and site-specific neoantigen-reactive clones were observed, demonstrating that a polyclonal TCR repertoire associated with a single neoantigen can vary by site. Although under-sampling or lack of detection of TCR sequences with low abundance may have contributed to this finding, such site-specific T-cell clone persistence has been observed in healthy individuals, notably for tissue-resident memory T-cells^(53, 54)^. In cancer, while site-specific TCR repertoires across metastases are often attributed to differential neoantigen expression and/or presentation ^(55)^, our data would suggest a possible additional factor for this heterogeneity.

The disappointing results of immunotherapy clinical trials in PNET^(27)^ show that ICB alone may be insufficient to induce clinically relevant anti-tumor immune responses in these patients. Combination therapy with ICB and NDT may provide additional benefit, as reported for other malignancies^(9, 10)^. However, data exploring NDT in patients with PNETs are scarce. Notably, Li *et al.*^(56)^ reported evidence of T-cell response to a multiepitope DNA vaccine in one patient with metastatic PNET and a clinical trial (NCT03412877) is currently investigating neoantigen-reactive adoptive T-cell therapy in patients with metastatic cancer, including PNETs. The identification of neoantigen-reactive TILs in our study supports further exploration of NDT in patients with PNETs. The clinical success of adjuvant vaccination in the minimal residual disease setting^(6, 7)^ suggests that vaccination after the clearance of macroscopic disease by surgery is promising.

Our findings also underscore the potential of ubiquitous neoantigens as appropriate targets for patients with metastatic PNETs. This neoantigen selection strategy could be applied to other solid tumors with advanced metastatic disease, including low TMB tumors, where access to both primary and metastatic sites is clinically possible. Neoantigen quality is crucial for designing NDT and current assessment typically considers factors such as expression, MHC binding, dissimilarity with self-antigens, clonality, and T-cell recognition^(57, 58, 59)^. However, the sharing status of targeted mutations across metastatic sites is often overlooked. We propose that this sharing status of the targeted mutations, regardless of their clonal nature, should be considered a key criterion in evaluating neoantigen quality and designing NDT for metastatic disease patients.

Practical challenges in implementing this approach include accessing samples from multiple metastatic sites; while this may be possible for tumors like PNETs, it may be more difficult for other cancers. The use of liquid biopsies could mitigate this difficulty by providing a less invasive method to detect mutations most likely to be shared across metastases^(60)^. Although our study shows promising results, the role of ubiquitous neoantigen-reactive T-cells in patients with metastatic disease requires confirmation in larger cohorts. Research autopsy studies may provide further insights into this concept, offering a comprehensive view of neoantigen distribution and T-cell responses across metastatic sites^(61)^.

In conclusion, our study provides the first evidence of pre-existing ubiquitous neoantigen-reactive TILs across several metastatic sites in a patient with PNET. These findings support the development of NDT in patients with PNET and suggest that mutations shared across multiple tumor sites can be immunogenic and lead to expansion of associated neoantigen-reactive TILs. This discovery not only has implications for PNET treatment but may also inform therapeutic strategies for other metastatic solid tumors.

## DECLARATIONS

### Ethics approval and consent to participate

Ethical approval was obtained from the local research ethics committee, LREC reference 14-SC-0186 150975 (2016) and 17/WA/0241 (2018) and written informed consent was provided by the patient.

### Patient consent for publication

Not applicable

### Availability of data and material

All data relevant to the study are included in the main manuscript or supplemental material. Bulk RNA and whole exome sequencing data have been deposited at the European Genome-phenome Archive (EGA), hosted by the European Bioinformatics Institute and the Centre for Genomic Regulation, under accession number EGAS00001006722. Proteomics data have been deposited in the ProteomeXchange Consortium via the PRIDE partner repository with the identifiers PXD037449 and 10.6019/PXD037449. Single-cell RNA/TCR/CITE and bulk TCR sequencing data can be accessed in Gene Expression Omnibus (GEO) under accession number GSE301291. All codes for bioinformatic analysis were deposited in our GitHub repository (https://github.com/vijaybioinfo/Metastatic-Neoantigen-Tcells-PNET)

## Acknowledgments

We thank Maria Lopez and the Research Pathology core at Southampton University Hospitals for performing immunohistochemistry on the paraffin-embedded material and James Pickering for QuPath analysis of cell markers. We thank Benjamin Johnson and the clinical trial assistant team at Southampton University Hospitals for sample collection. We thank Oliver Wood, Divya Singh, Konstantinos Boukas and the flow cytometry core from the La Jolla Institute of Immunology for technical support.

## Authors’ contributions

J-BT, AB: experimental work, bioinformatic evaluation, data interpretation, paper writing, paper review; KM: experimental work, data interpretation, paper writing, paper review; FECa, JT, GS: bioinformatic evaluation, data interpretation, paper review; PSi, ECu, HS, BN: experimental work, data interpretation, paper review; AT: sample collection; data interpretation, paper review; SB-R: histological assessment, data interpretation, paper review; LC, JC, SPS, PSk, PV, NS: data interpretation, paper review; CO: conceived, supervised and led the work, data interpretation, paper writing, paper review.

## Abbreviations

NDT: Neoantigen-directed therapy
TMB: Tumor mutational burden
PNET: Pancreatic neuroendocrine tumor
TIL: Tumor-infiltrating T-cell
ICB: Immune checkpoint blockade
CK: Cytokeratin, e.g. CK7, CK20
FFPE: Formalin-fixed, paraffin-embedded
FOLFOX: Fluorouracil, leucovorin calcium and oxaliplatin
IHC: Immunohistochemistry
MHC-I/MHC-II: Major Histocompatibility Complex Class I and Class II
EGA: European Genome-Phenome Archive
RNAseq: RNA sequencing
TPM: Transcripts per million
GEO: Gene Expression Omnibus
HNSCC: Head and neck squamous cell carcinoma
WES: Whole exome sequencing
SNV: Single nucleotide variant
ELISpot: Enzyme-linked immunosorbent spot assay
LP: Long peptide (20mer, with mutation at position 6 or 15)
wt: Wild-type
HLA: Human Leukocyte Antigen
IEDB: Immune Epitope Database
ME: Minimal epitope
PBMC: Peripheral blood mononuclear cell
IL: Interleukin, e.g. IL-2, IL-7, IL-15
TCRseq: T-cell receptor sequencing
PHA: Phytohemagglutinin
SFC: Spot forming cells
FBS: Fetal bovine serum
UMI: Unique molecular identifier
PBS: Phosphate buffered saline
UMAP: Uniform manifold approximation and projection
ADT: Antibody-derived tag
DP: Double-negative
DN: Double-positive
GEX: Gene expression
LC-MS/MS: Liquid chromatography-tandem mass spectrometry
FDR: False discovery rate
CD8act: Activated CD8 T-cell
CD4act: Activated CD4 T-cell
T_CM_: Central memory T-cell
T_EM_: Effector memory T-cell

---

**Supplemental Figure S1.**
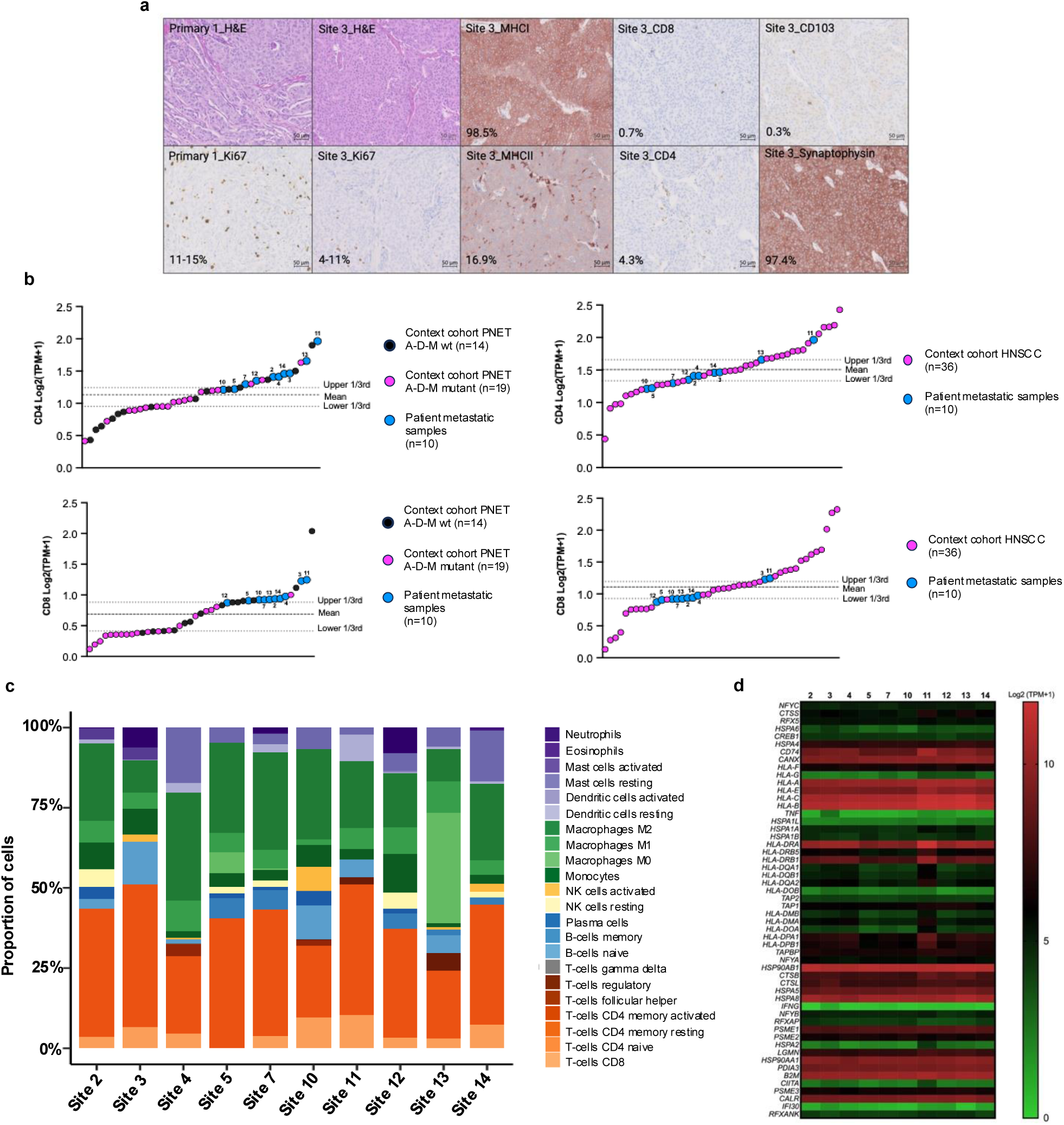
Tumor characterization |. **a)** Representative images for hematoxylin and eosin (H&E) and immunohistochemical staining of the primary tumor (Ki67 only) and one representative metastatic site (site 3) for Ki-67, MHC-I, MHC-II, CD4, CD8, CD103 and synaptophysin. **b)** Scatter plot showing gene transcripts expressed in Log2 normalized TPM for CD4 (top panels) and CD8 (bottom panels) obtained with whole tumor RNAseq of metastatic sites (n=10, blue circles) by comparison with contextualization cohorts: a cohort (n=33) of A-D-M mutant (pink circles, n=19) and A-D-M wild type (black circles, n =14) PNET (left panels)^28^ and a cohort of head and neck squamous cell carcinoma (HNSCC) (pink circles, n = 36)^29^ (right panels). Dashed lines represent the mean, dotted lines represent upper and lower 3rd percentiles. **c)** Bar plot showing the proportions of 22 immune cell subsets in metastatic sites (n=10), generated using CIBERSORT. **d)** Heatmap showing the scaled (Log2 (TPM+1)) expression of 54 genes involved antigen processing and presentation by MHC-I and MHC-II across metastatic sites (n=10).

**Supplemental Figure S2.**
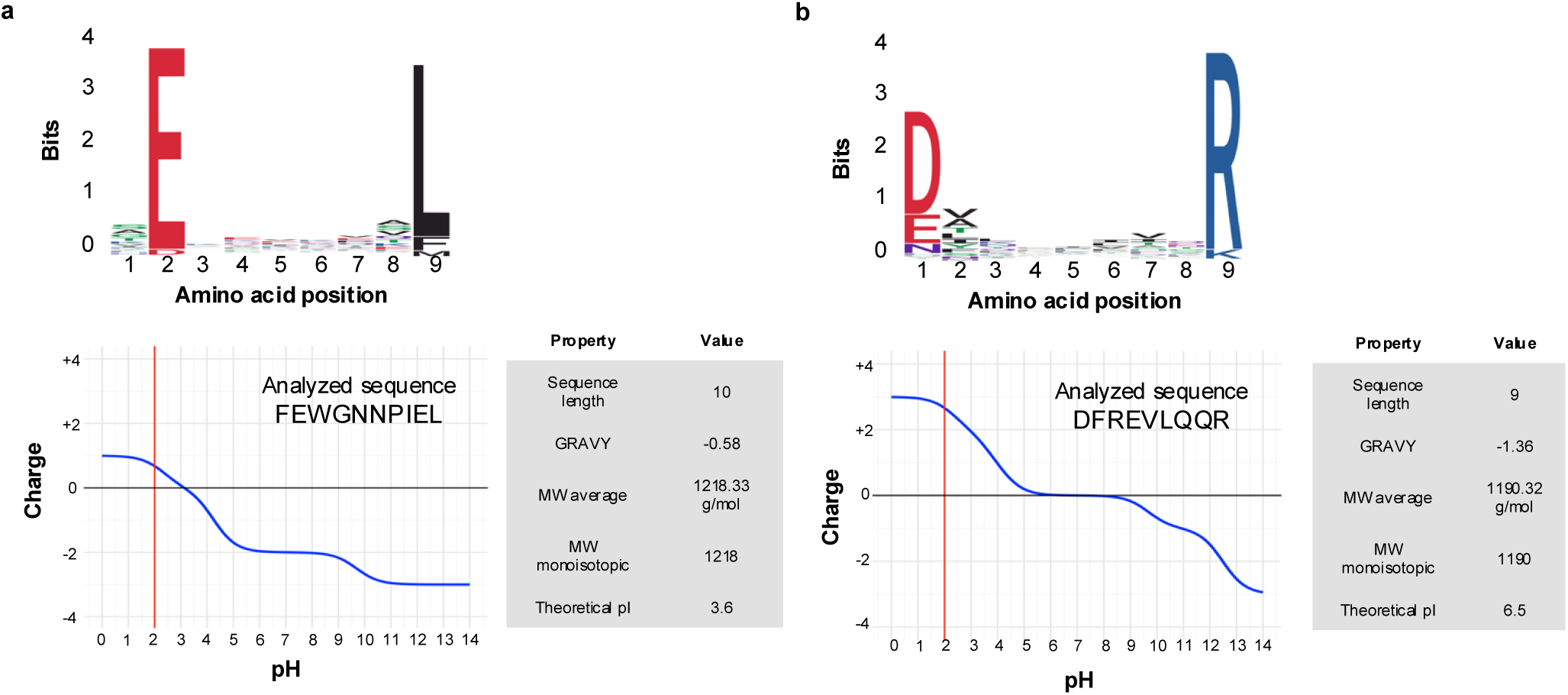
Immunopeptidomics |. **a)** HLA-B*40:02 NPTX2-derived peptides. Sequence binding motif of all peptides eluted from HLA-B*40:02 (n=9) (top panel). The bit score (bits, y axis) indicates the residue content for each position in the sequence: polar amino acids (green; G, S, T, Y, C, Q, N), basic amino acids (blue; K, R, H), acidic amino acids (red; D, E) and hydrophobic amino acids (black; A, V, L, I, P, W, F, M). Hydrophobicity score and net electrical charge (bottom left panel) for the predicted mNPTX2-derived peptide (FEWGNNPIEL), not observed in LC-MS/MS. The blue line denotes the change in peptide charge with varying pH; the red line indicates the peptide charge at pH2, utilized for MS. The table (bottom right panel) indicates the properties of the predicted mNPTX2-derived peptide. **b)** HLA-A*33:01 wtNPTX2-derived peptides. Sequence binding motif of all peptides eluted from HLA-A*33:01 (n=9) (top panel). Hydrophobicity score and net electrical charge (bottom left panel) for the observed wtNPTX2-derived peptide (DFREVLQQR) captured in LC-MS/MS. The table (bottom right panel) indicates the properties of the observed wtNPTX2-derived peptide.

**Supplemental Figure S3.**
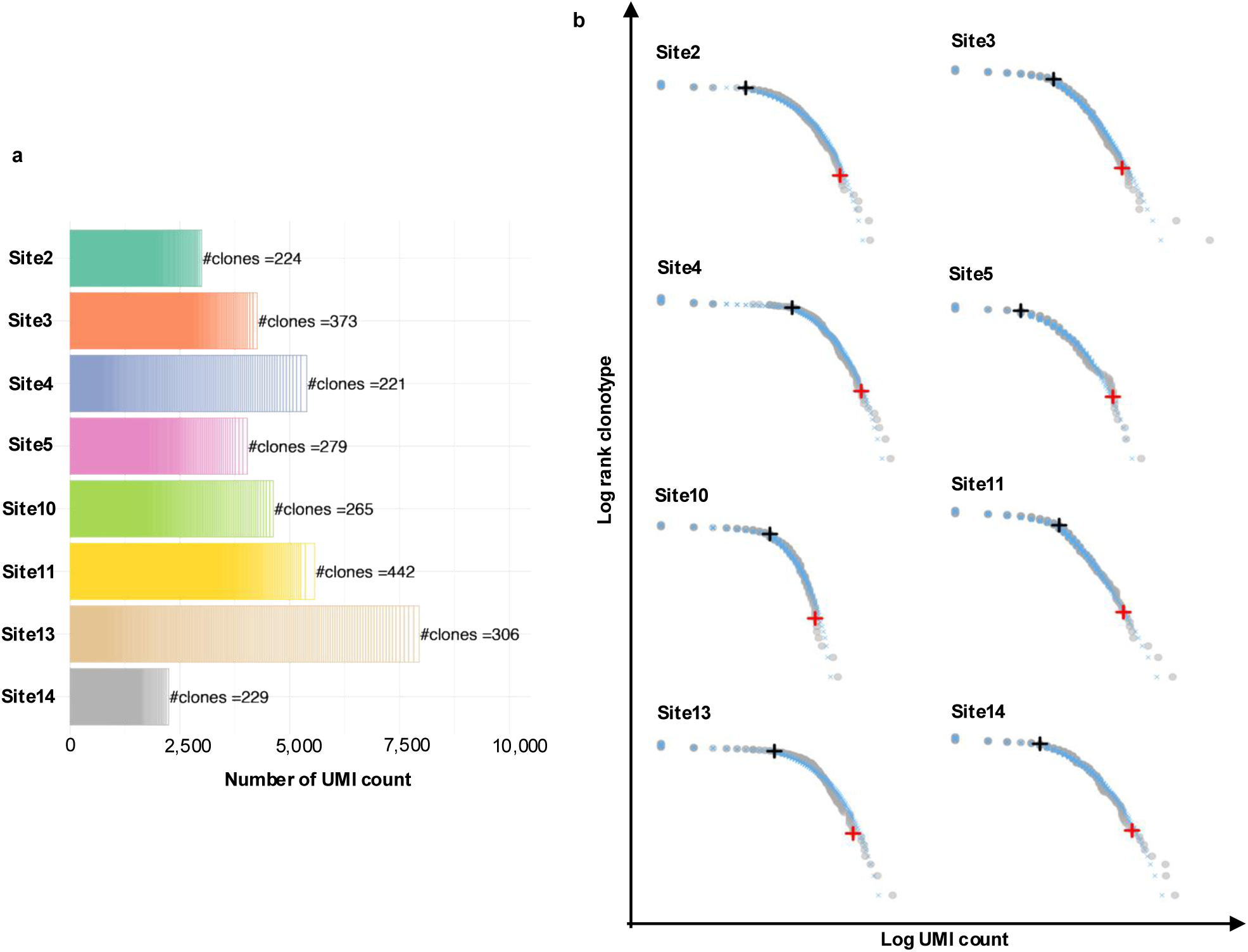
Clonotype analysis in metastatic sites |. **a)** UMI count distribution and clonotype richness of unique TCRβ clonotype across metastatic sites (n=8). **b)** Dot plots showing the distribution of UMI count (log UMI count) for each ranked (log rank) clonotype across all metastatic sites (n=8). The grey dots represent the actual dataset, and the blue crosses represent the theoretical UMI count for each clone, fitted using the discrete Gamma-GPD spliced threshold model. The black and red crosses represent the thresholds associated with “expanded” and “highly expanded” clonotypes, respectively.

**Supplemental Figure S4.**
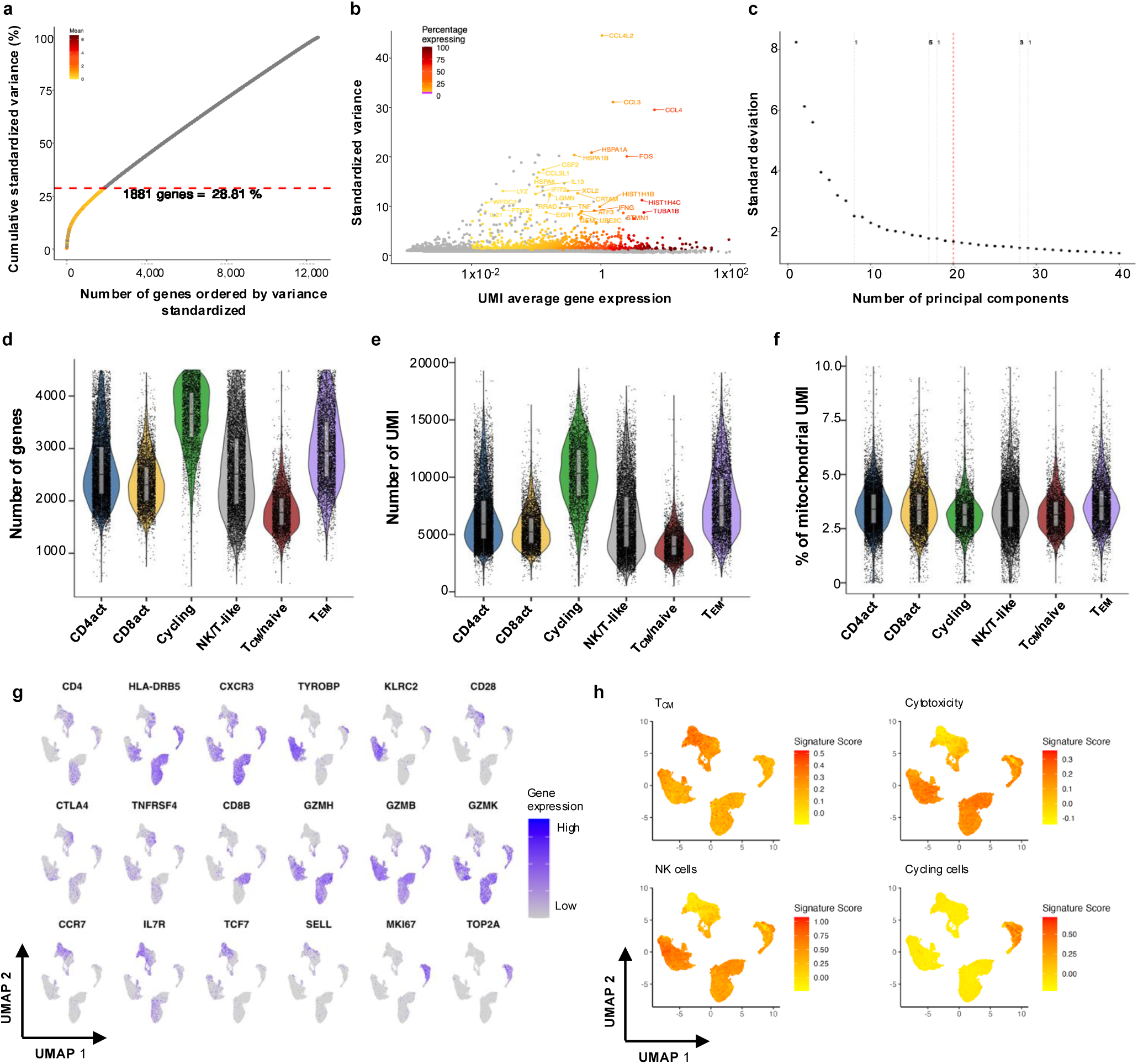
Single-cell transcriptomics: quality control and cluster assignment |. **a)** Scatter plot showing the cumulative standardized variance for genes (n=17,804) expressed in more than 0.1% of the cells and with a mean UMI >0.01 (y axis) in function of the number of genes, ranked from highest to lowest standardized variance values (x axis). Genes below the red dotted line were selected (n=1,881) and are part of the highest variable genes explaining 28.8% of the cumulative standardized variance. Genes are colored based on the UMI mean expression obtained with Seurat. **b)** Scatter plot showing the distribution of genes ordered based on their UMI mean expression values (x axis) in function of their individual standardized variance values (y axis). Genes colored in red-orange are the 1,881 highest variable genes with a UMI mean expression value >0.01. **c)** Scatter plot showing the standardized variation for first 40 principal components (PCs) using the 1,773 most variable genes. Red lines indicate the number of PCs selected for clustering analysis with the Seurat methodology. **d-f)** scRNAseq quality control: Violin plots showing the per-cell distribution of (**d**) unique genes, (**e**) UMI and (**f**) percentage of UMI mapped to mitochondrial genome in 35,178 T-cells across clusters in mNPTX2.LP and mNPTX2.ME expanded T-cells. Box plots extend from the 25^th^ to 75^th^ percentile and the center line represents the median. Whiskers are bounded by 25^th^ percentile - 1.5*interquartile range and 75^th^ percentile + 1.5*interquartile range. **g)** UMAPs showing expression (log normalized count) of canonical genes across cell clusters. **h)** UMAPs showing gene set enrichment score for T-cell memory, cytotoxicity, natural killer (NK) cells and cycling cells signatures.

**Supplemental Figure S5.**
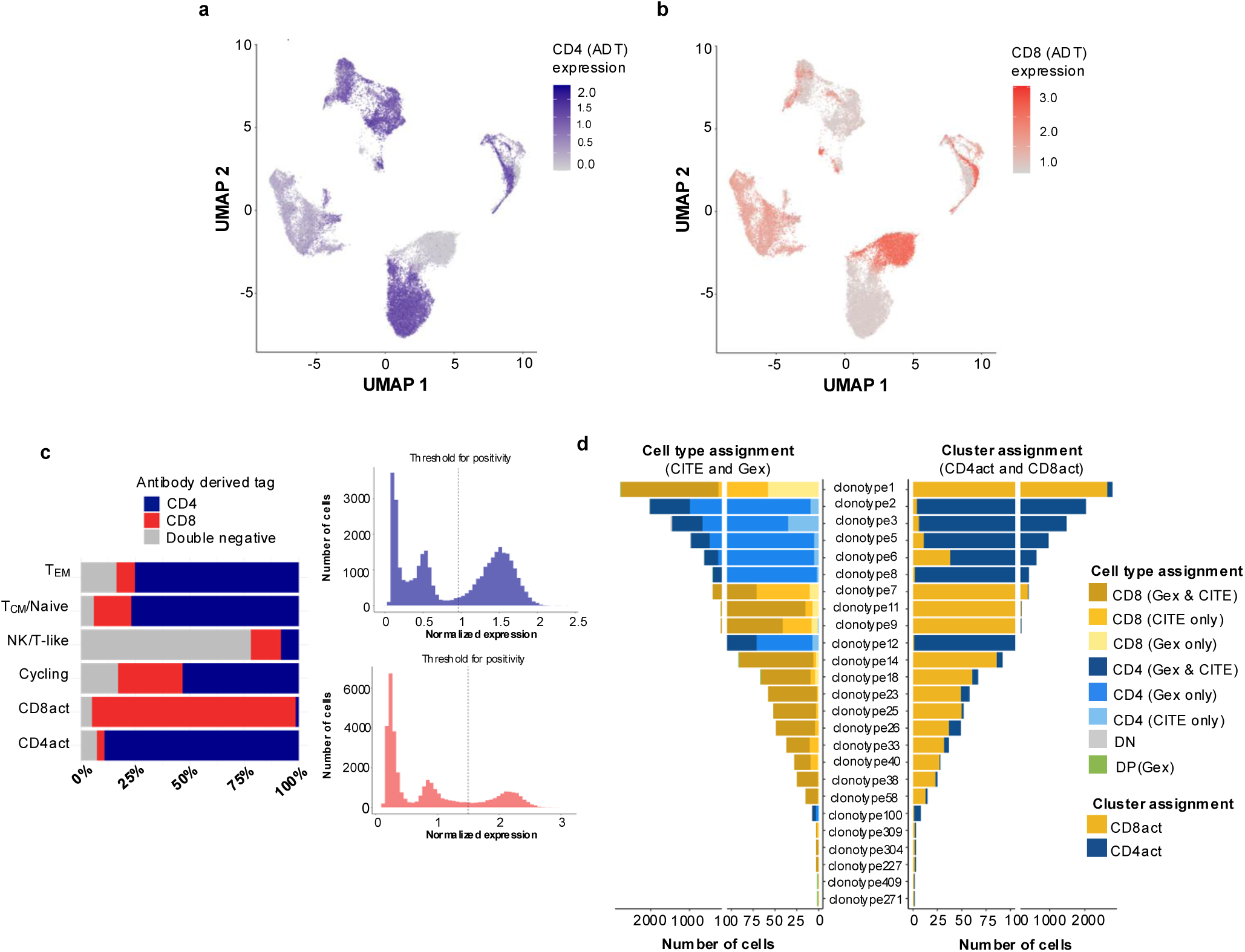
Single-cell transcriptomics: cell type assignment |. **a)** UMAP showing the expression of CD4 ADT. **b)** UMAP showing the expression of CD8 ADT. **c)** Bar charts (left panel) showing the proportion of CD4-ADT positive, CD8-ADT positive and double negative cells per cluster. Histograms (right panel) showing the distribution of CD4-ADT (top panel) and CD8-ADT (bottom panel) expression levels across all cells. The vertical bar shows the threshold used for a positive cell. **d**) Bar plot (left panel) showing the proportion of cells defined as CD4^+^ or CD8^+^ based on CITEseq and gene expression data for each clone shared by both CD4act and CD8act clusters. Bar plot (right panel) showing the proportion of cells in CD4act and CD8act for each clone projected in both clusters.

**Supplemental Figure S6.**
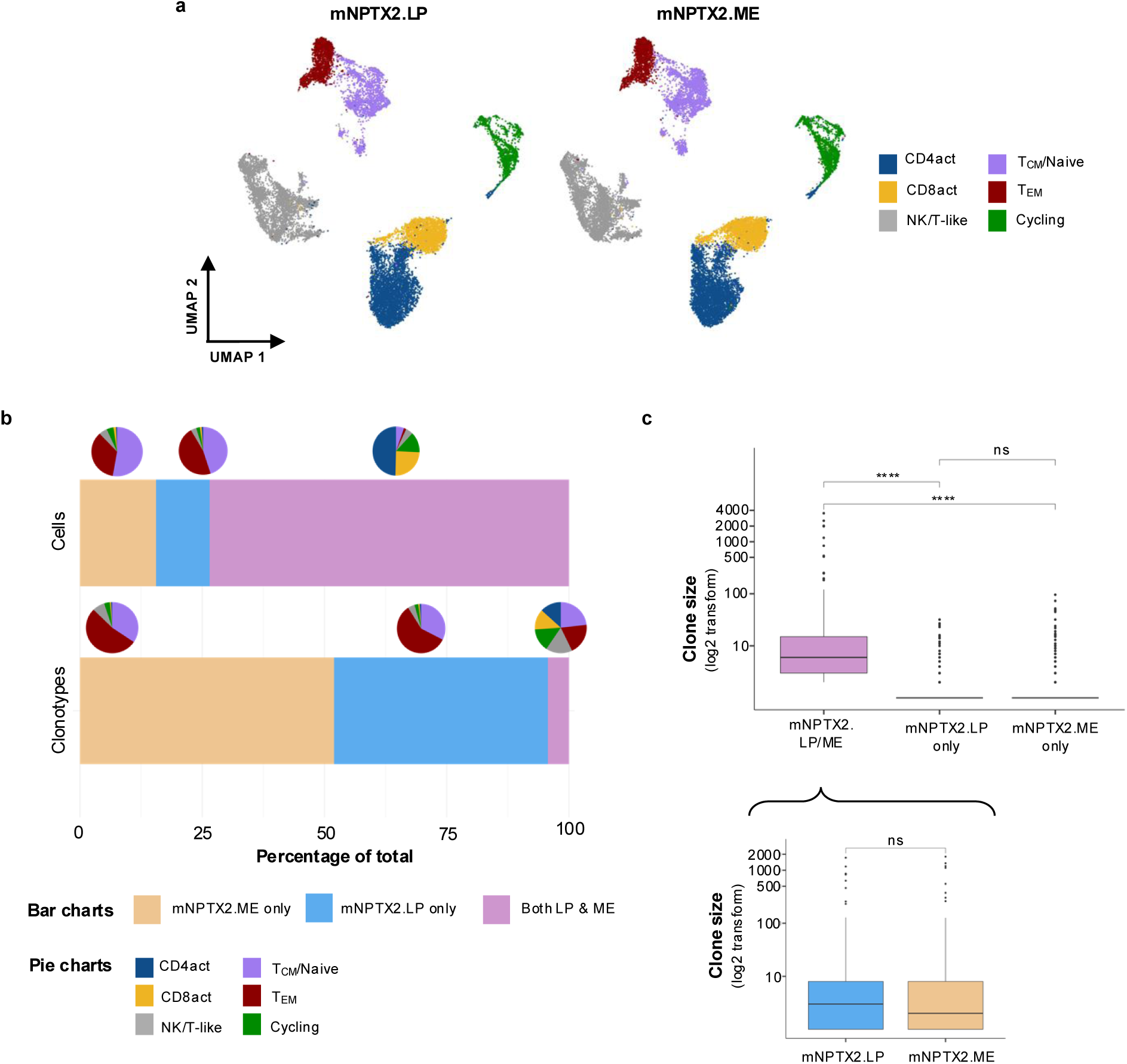
Single-cell transcriptomics: comparison across peptide stimulation conditions |. **a)** UMAPs showing cells originating from mNPTX2.LP (left panel) and mNPTX2.ME (right panel) expanded T-cells. **b)** Bar plots showing the proportion of cells (top panel) and clonotypes (bottom panel) associated with mNPTX2.LP and mNPTX2.ME stimulation. The pie charts show the associated distribution of cluster for each clonotype/cell associated with the two peptide stimulation conditions. **c)** Box plot showing the clone size (log2 scale) for each T-cell clone present in mNPTX2.LP only, mNPTX2.ME only or both datasets. Within the clones present in both datasets, a box plot (bottom panel) shows the clone size for each clone for each peptide stimulation. Statistical significance between peptide stimulation was assessed using the two-sided Wilcoxon test with p-values adjusted for multiple comparisons using the Benjamini-Hochberg method. **** indicates a p-value <0.0001.

**Supplemental Figure S7.**
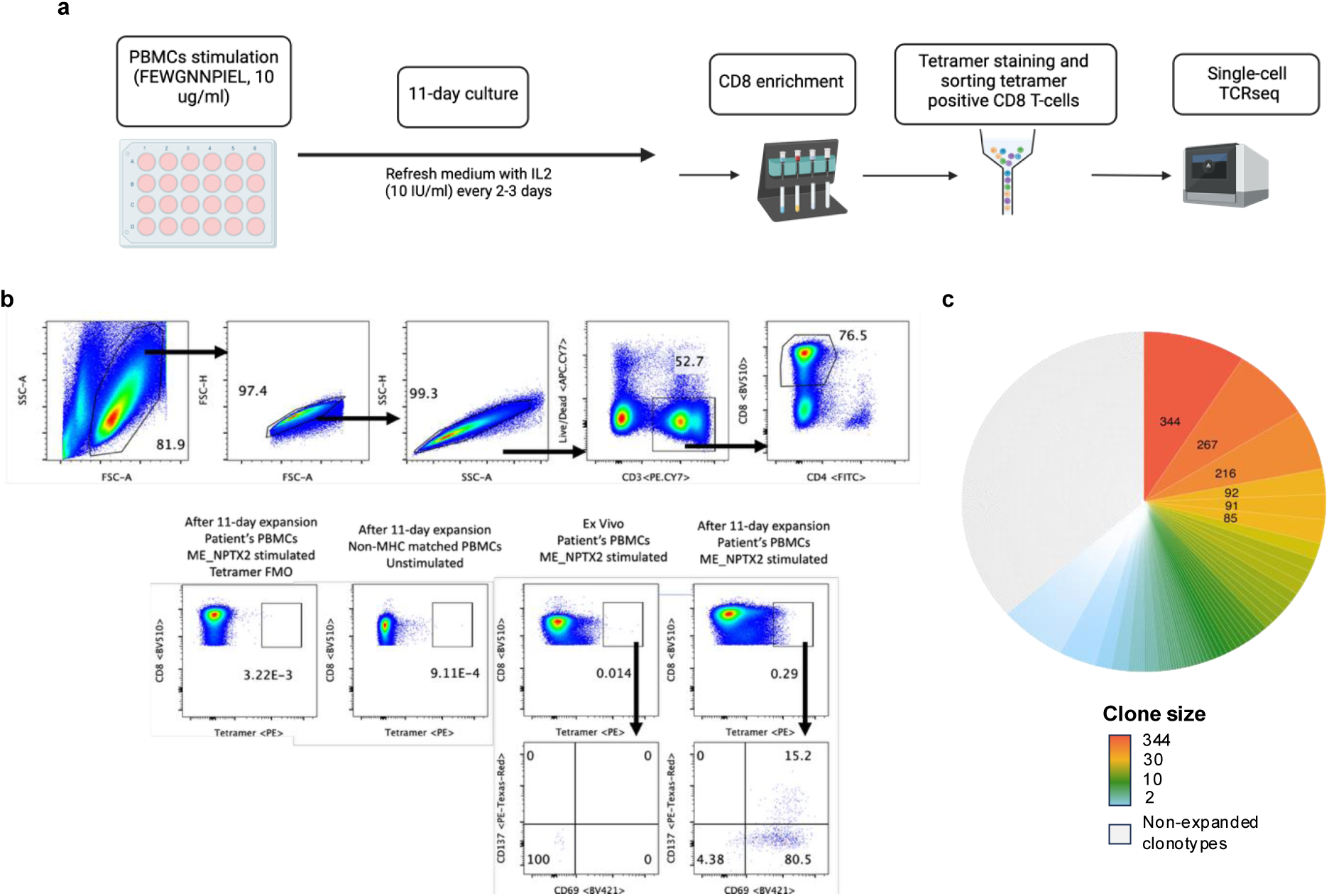
Tetramer data |. **a)** Schematic showing the workflow used to identify NPTX2-reactive CD8 T-cell clonotypes utilizing an HLA-B*40:02-restriced mNPTX2.ME.HLA-I (FEWGNNPIEL) tetramer. **b)** Gating strategy used for the tetramer data. **c)** Pie chart showing the distribution of clone size from tetramer-positive CD8 T-cells clones.

**Supplemental Figure S8.**
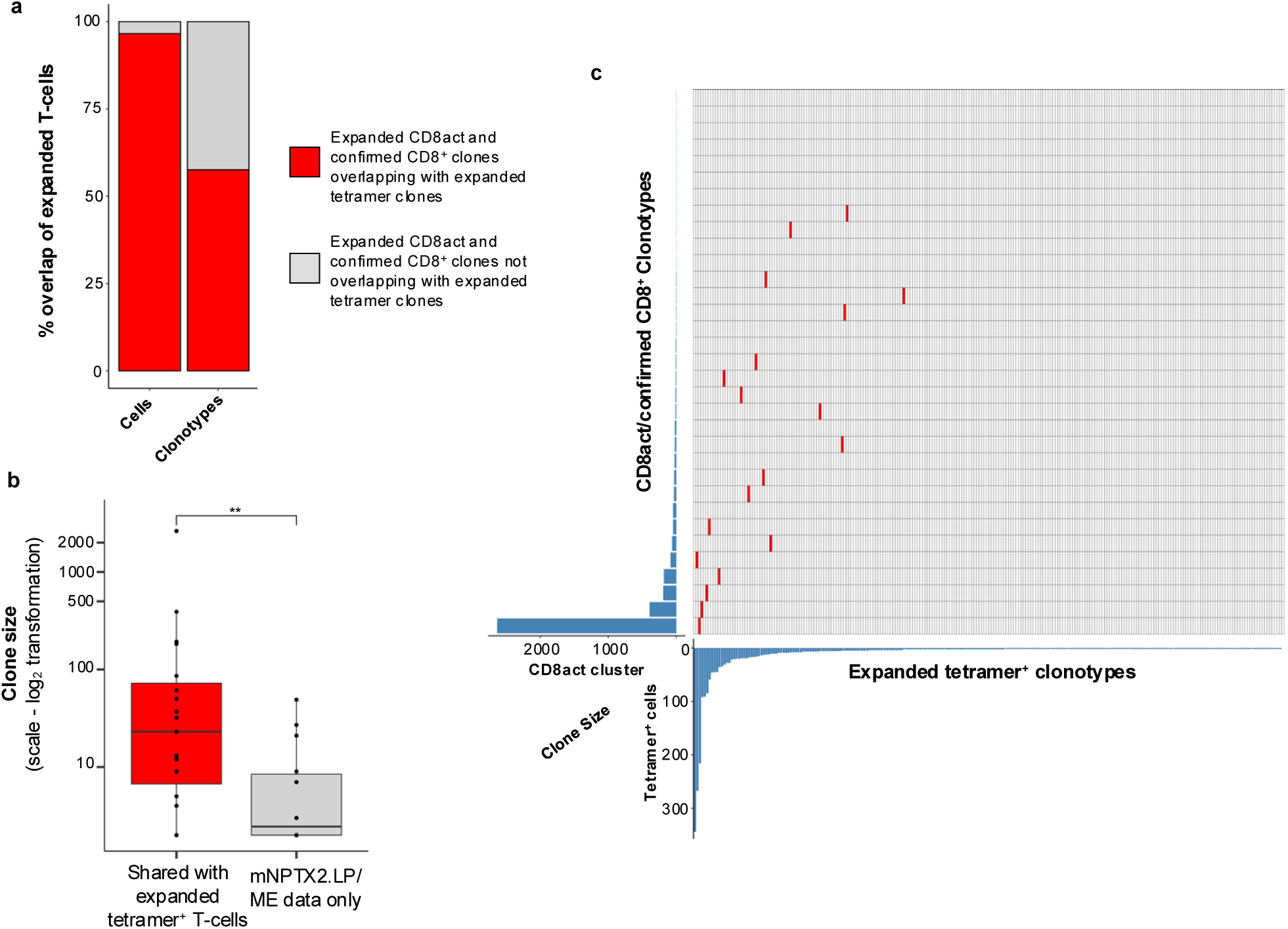
Overlap between tetramer and mNPTX2.LP/ME expanded T-cells |. **a)** Bar chart showing the proportion of clonotypes and cells overlapping between tetramer data and CD8act CD8^+^ clones from mNPTX2.LP/ME expanded T-cells. **b)** Box plot comparing the clone size for CD8act CD8^+^ clones shared between tetramer data and CD8act CD8^+^ clones from mNPTX2.LP/ME expanded T-cells. p-value derived from two-sided Wilcoxon test (p-value = 0.03). **c)** Heatmap showing the clonotype overlap (in red) between CD8act CD8^+^ clones from mNPTX2.LP/ME expanded T-cells (y axis) and tetramer-positive CD8 T-cells (x axis). The bar chart represents the clone size for each clone in the two datasets. Only expanded T-cells (clone size ≥ 2) are shown.

## References

1. Esposito M, Ganesan S, Kang Y. Emerging strategies for treating metastasis. Nat Cancer. 2021;2(3):258–70.

2. Ganesh K, Massague J. Targeting metastatic cancer. Nat Med. 2021;27(1):34–44.

3. Biswas N, Chakrabarti S, Padul V, Jones LD, Ashili S. Designing neoantigen cancer vaccines, trials, and outcomes. Front Immunol. 2023;14:1105420.

4. Borden ES, Buetow KH, Wilson MA, Hastings KT. Cancer Neoantigens: Challenges and Future Directions for Prediction, Prioritization, and Validation. Front Oncol. 2022;12:836821.

5. Schumacher TN, Schreiber RD. Neoantigens in cancer immunotherapy. Science. 2015;348(6230):69–74.

6. Braun DA, Moranzoni G, Chea V, McGregor BA, Blass E, Tu CR, et al. A neoantigen vaccine generates antitumour immunity in renal cell carcinoma. Nature. 2025;639(8054):474–82.

7. Ottensmeier CHH, Delord J-P, Lalanne A, Lantz O, Jamet C, Tavernaro A, et al. Safety and immunogenicity of TG4050: A personalized cancer vaccine in head and neck carcinoma. Journal of Clinical Oncology. 2023;41(16_suppl):6082.

8. Katsikis PD, Ishii KJ, Schliehe C. Challenges in developing personalized neoantigen cancer vaccines. Nat Rev Immunol. 2024;24(3):213–27.

9. Linette GP, Becker-Hapak M, Skidmore ZL, Baroja ML, Xu C, Hundal J, et al. Immunological ignorance is an enabling feature of the oligo-clonal T cell response to melanoma neoantigens. Proc Natl Acad Sci U S A. 2019;116(47):23662–70.

10. Rappaport AR, Kyi C, Lane M, Hart MG, Johnson ML, Henick BS, et al. A shared neoantigen vaccine combined with immune checkpoint blockade for advanced metastatic solid tumors: phase 1 trial interim results. Nat Med. 2024;30(4):1013–22.

11. Angelova M, Mlecnik B, Vasaturo A, Bindea G, Fredriksen T, Lafontaine L, et al. Evolution of Metastases in Space and Time under Immune Selection. Cell. 2018;175(3):751–65 e16.

12. De Mattos-Arruda L, Sammut SJ, Ross EM, Bashford-Rogers R, Greenstein E, Markus H, et al. The Genomic and Immune Landscapes of Lethal Metastatic Breast Cancer. Cell Rep. 2019;27(9):2690–708 e10.

13. Reiter JG, Makohon-Moore AP, Gerold JM, Heyde A, Attiyeh MA, Kohutek ZA, et al. Minimal functional driver gene heterogeneity among untreated metastases. Science. 2018;361(6406):1033–7.

14. Chen R, Li J, Fujimoto J, Hong L, Hu X, Quek K, et al. Immunogenomic intertumor heterogeneity across primary and metastatic sites in a patient with lung adenocarcinoma. J Exp Clin Cancer Res. 2022;41(1):172.

15. Gerlinger M, Rowan AJ, Horswell S, Math M, Larkin J, Endesfelder D, et al. Intratumor heterogeneity and branched evolution revealed by multiregion sequencing. N Engl J Med. 2012;366(10):883–92.

16. Hiam-Galvez KJ, Allen BM, Spitzer MH. Systemic immunity in cancer. Nat Rev Cancer. 2021;21(6):345–59.

17. Chen Z, Zhang S, Han N, Jiang J, Xu Y, Ma D, et al. A Neoantigen-Based Peptide Vaccine for Patients With Advanced Pancreatic Cancer Refractory to Standard Treatment. Front Immunol. 2021;12:691605.

18. D’Alise AM, Leoni G, Cotugno G, Siani L, Vitale R, Ruzza V, et al. Phase I Trial of Viral Vector-Based Personalized Vaccination Elicits Robust Neoantigen-Specific Antitumor T-Cell Responses. Clin Cancer Res. 2024;30(11):2412–23.

19. Leidner R, Sanjuan Silva N, Huang H, Sprott D, Zheng C, Shih YP, et al. Neoantigen T-Cell Receptor Gene Therapy in Pancreatic Cancer. N Engl J Med. 2022;386(22):2112–9.

20. Mork SK, Skadborg SK, Albieri B, Draghi A, Bol K, Kadivar M, et al. Dose escalation study of a personalized peptide-based neoantigen vaccine (EVX-01) in patients with metastatic melanoma. J Immunother Cancer. 2024;12(5).

21. Sahin U, Derhovanessian E, Miller M, Kloke BP, Simon P, Lower M, et al. Personalized RNA mutanome vaccines mobilize poly-specific therapeutic immunity against cancer. Nature. 2017;547(7662):222–6.

22. Yarchoan M, Gane EJ, Marron TU, Perales-Linares R, Yan J, Cooch N, et al. Personalized neoantigen vaccine and pembrolizumab in advanced hepatocellular carcinoma: a phase 1/2 trial. Nat Med. 2024;30(4):1044–53.

23. Parkhurst M, Goff SL, Lowery FJ, Beyer RK, Halas H, Robbins PF, et al. Adoptive transfer of personalized neoantigen-reactive TCR-transduced T cells in metastatic colorectal cancer: phase 2 trial interim results. Nat Med. 2024;30(9):2586–95.

24. McGranahan N, Furness AJ, Rosenthal R, Ramskov S, Lyngaa R, Saini SK, et al. Clonal neoantigens elicit T cell immunoreactivity and sensitivity to immune checkpoint blockade. Science. 2016;351(6280):1463–9.

25. Franko J, Feng W, Yip L, Genovese E, Moser AJ. Non-functional neuroendocrine carcinoma of the pancreas: incidence, tumor biology, and outcomes in 2,158 patients. J Gastrointest Surg. 2010;14(3):541–8.

26. Priestley P, Baber J, Lolkema MP, Steeghs N, de Bruijn E, Shale C, et al. Pan-cancer whole- genome analyses of metastatic solid tumours. Nature. 2019;575(7781):210–6.

27. Garcia-Torralba E, Garcia-Lorenzo E, Doger B, Spada F, Lamarca A. Immunotherapy in Neuroendocrine Neoplasms: A Diamond to Cut. Cancers (Basel). 2024;16(14).

28. Chan CS, Laddha SV, Lewis PW, Koletsky MS, Robzyk K, Da Silva E, et al. ATRX, DAXX or MEN1 mutant pancreatic neuroendocrine tumors are a distinct alpha-cell signature subgroup. Nat Commun. 2018;9(1):4158.

29. Wood O, Woo J, Seumois G, Savelyeva N, McCann KJ, Singh D, et al. Gene expression analysis of TIL rich HPV-driven head and neck tumors reveals a distinct B-cell signature when compared to HPV independent tumors. Oncotarget. 2016;7(35):56781–97.

30. Koch H, Starenki D, Cooper SJ, Myers RM, Li Q. powerTCR: A model-based approach to comparative analysis of the clone size distribution of the T cell receptor repertoire. PLoS Comput Biol. 2018;14(11):e1006571.

31. Stuart T, Butler A, Hoffman P, Hafemeister C, Papalexi E, Mauck WM, 3rd, et al. Comprehensive Integration of Single-Cell Data. Cell. 2019;177(7):1888–902 e21.

32. Denisenko E, Guo BB, Jones M, Hou R, de Kock L, Lassmann T, et al. Systematic assessment of tissue dissociation and storage biases in single-cell and single-nucleus RNA-seq workflows. Genome Biol. 2020;21(1):130.

33. van den Brink SC, Sage F, Vertesy A, Spanjaard B, Peterson-Maduro J, Baron CS, et al. Single-cell sequencing reveals dissociation-induced gene expression in tissue subpopulations. Nat Methods. 2017;14(10):935–6.

34. Meckiff BJ, Ramirez-Suastegui C, Fajardo V, Chee SJ, Kusnadi A, Simon H, et al. Imbalance of Regulatory and Cytotoxic SARS-CoV-2-Reactive CD4(+) T Cells in COVID-19. Cell. 2020;183(5):1340–53 e16.

35. Patil VS, Madrigal A, Schmiedel BJ, Clarke J, O’Rourke P, de Silva AD, et al. Precursors of human CD4(+) cytotoxic T lymphocytes identified by single-cell transcriptome analysis. Sci Immunol. 2018;3(19).

36. Pauken KE, Lagattuta KA, Lu BY, Lucca LE, Daud AI, Hafler DA, et al. TCR-sequencing in cancer and autoimmunity: barcodes and beyond. Trends Immunol. 2022;43(3):180–94.

37. Szabo PA, Levitin HM, Miron M, Snyder ME, Senda T, Yuan J, et al. Single-cell transcriptomics of human T cells reveals tissue and activation signatures in health and disease. Nat Commun. 2019;10(1):4706.

38. Terekhova M, Swain A, Bohacova P, Aladyeva E, Arthur L, Laha A, et al. Single-cell atlas of healthy human blood unveils age-related loss of NKG2C(+)GZMB(-)CD8(+) memory T cells and accumulation of type 2 memory T cells. Immunity. 2023;56(12):2836–54 e9.

39. Nicholas B, Bailey A, McCann KJ, Wood O, Walker RC, Parker R, et al. Identification of neoantigens in oesophageal adenocarcinoma. Immunology. 2023;168(3):420–31.

40. Perez-Riverol Y, Bai J, Bandla C, Garcia-Seisdedos D, Hewapathirana S, Kamatchinathan S, et al. The PRIDE database resources in 2022: a hub for mass spectrometry-based proteomics evidences. Nucleic Acids Res. 2022;50(D1):D543–D52.

41. Alexandrov LB, Kim J, Haradhvala NJ, Huang MN, Tian Ng AW, Wu Y, et al. The repertoire of mutational signatures in human cancer. Nature. 2020;578(7793):94–101.

42. Declercq A, Bouwmeester R, Hirschler A, Carapito C, Degroeve S, Martens L, Gabriels R. MS(2)Rescore: Data-Driven Rescoring Dramatically Boosts Immunopeptide Identification Rates. Mol Cell Proteomics. 2022;21(8):100266.

43. Timp W, Timp G. Beyond mass spectrometry, the next step in proteomics. Sci Adv. 2020;6(2):8978.

44. Jiang Y, Dong YH, Zhao SW, Liu DY, Zhang JY, Xu XY, et al. Multiregion WES of metastatic pancreatic neuroendocrine tumors revealed heterogeneity in genomic alterations, immune microenvironment and evolutionary patterns. Cell Commun Signal. 2024;22(1):164.

45. Rogiers A, Lobon I, Spain L, Turajlic S. The Genetic Evolution of Metastasis. Cancer Res. 2022;82(10):1849–57.

46. Keskin DB, Anandappa AJ, Sun J, Tirosh I, Mathewson ND, Li S, et al. Neoantigen vaccine generates intratumoral T cell responses in phase Ib glioblastoma trial. Nature. 2019;565(7738):234–9.

47. Veatch JR, Lee SM, Shasha C, Singhi N, Szeto JL, Moshiri AS, et al. Neoantigen-specific CD4(+) T cells in human melanoma have diverse differentiation states and correlate with CD8(+) T cell, macrophage, and B cell function. Cancer Cell. 2022;40(4):393–409.e9.

48. Wu D, Gallagher DT, Gowthaman R, Pierce BG, Mariuzza RA. Structural basis for oligoclonal T cell recognition of a shared p53 cancer neoantigen. Nature Communications. 2020;11(1).

49. Sewell AK. Why must T cells be cross-reactive? Nat Rev Immunol. 2012;12(9):669–77.

50. Balachandran VP, Luksza M, Zhao JN, Makarov V, Moral JA, Remark R, et al. Identification of unique neoantigen qualities in long-term survivors of pancreatic cancer. Nature. 2017;551(7681):512–6.

51. Cavalluzzo B, Viuff MC, Tvingsholm SA, Ragone C, Manolio C, Mauriello A, et al. Cross- reactive CD8(+) T cell responses to tumor-associated antigens (TAAs) and homologous microbiota- derived antigens (MoAs). J Exp Clin Cancer Res. 2024;43(1):87.

52. Al Bakir M, Reading JL, Gamble S, Rosenthal R, Uddin I, Rowan A, et al. Clonal driver neoantigen loss under EGFR TKI and immune selection pressures. Nature. 2025;639(8056):1052–9.

53. Miron M, Meng W, Rosenfeld AM, Dvorkin S, Poon MML, Lam N, et al. Maintenance of the human memory T cell repertoire by subset and tissue site. Genome Med. 2021;13(1):100.

54. Poon MML, Caron DP, Wang Z, Wells SB, Chen D, Meng W, et al. Tissue adaptation and clonal segregation of human memory T cells in barrier sites. Nat Immunol. 2023;24(2):309–19.

55. Joshi K, de Massy MR, Ismail M, Reading JL, Uddin I, Woolston A, et al. Spatial heterogeneity of the T cell receptor repertoire reflects the mutational landscape in lung cancer. Nat Med. 2019;25(10):1549–59.

56. Li L, Zhang X, Wang X, Kim SW, Herndon JM, Becker-Hapak MK, et al. Optimized polyepitope neoantigen DNA vaccines elicit neoantigen-specific immune responses in preclinical models and in clinical translation. Genome Med. 2021;13(1):56.

57. Lang F, Schrors B, Lower M, Tureci O, Sahin U. Identification of neoantigens for individualized therapeutic cancer vaccines. Nat Rev Drug Discov. 2022;21(4):261–82.

58. McGranahan N, Swanton C. Neoantigen quality, not quantity. Sci Transl Med. 2019;11(506).

59. Zhang J, Caruso FP, Sa JK, Justesen S, Nam D-H, Sims P, et al. The combination of neoantigen quality and T lymphocyte infiltrates identifies glioblastomas with the longest survival. Communications Biology. 2019;2(1).

60. Abbosh C, Frankell AM, Harrison T, Kisistok J, Garnett A, Johnson L, et al. Tracking early lung cancer metastatic dissemination in TRACERx using ctDNA. Nature. 2023;616(7957):553–62.

61. Iacobuzio-Donahue CA, Michael C, Baez P, Kappagantula R, Hooper JE, Hollman TJ. Cancer biology as revealed by the research autopsy. Nat Rev Cancer. 2019;19(12):686–97.

